# Engineered protein destabilization reverses intrinsic immune evasion for candidate vaccine pan-strain KSHV and SARS-CoV-2 antigens

**DOI:** 10.1101/2024.10.22.619692

**Authors:** Li Wan, Bingxian Xie, Masahiro Shuda, Greg Delgoffe, Yuan Chang, Patrick S. Moore

## Abstract

Both Kaposi sarcoma herpesvirus LANA and SARS coronavirus 2 RdRp/nsp12 are highly conserved replication proteins that evade immune processing. By deleting the LANA central repeat 1 domain (LANA^ΔCR1^) or by dividing RdRp into two separated fragments (RdRp^Frag^) to maximize nascent protein mis-folding, cis peptide presentation was increased. Native LANA or RdRp SIINFEKL fusion proteins expressed in MC38 cancer cells were not recognized by activated OT-1 CD8^+^ cells against SIINFEKL but cytotoxic recognition was restored by expression of the corresponding modified proteins. Immunocompetent syngeneic mice injected with LANA- or RdRp-SIINFEKL MC38 cells developed rapidly-growing tumors with short median survival times. Mice injected with LANA^ΔCR1^- or RdRp^Frag^-SIINFEKL had partial tumor regression, slower tumor growth, longer median survival, as well as increased effector-specific tumor-infiltrating lymphocytes. These mice developed robust T cell responses lasting at least 90 days post-injection that recognized native viral protein epitopes. Engineered vaccine candidate antigens can unmask virus-specific CTL responses that are typically suppressed during native viral infection.

## INTRODUCTION

Successful cytotoxic T cell lymphocyte (CTL) responses require immunologic priming in which foreign peptide antigens are presented to CD8^+^ T cells by MHC class I (MHC I) antigen presenting cells (APC) to initiate priming expansion of antigen-specific memory T cell clones (*1*). MHC I peptide APC presentation involves intracellular proteasomal proteolysis, either during nascent protein translation when misfolded proteins are detected (so-called defective ribosomal products (DRiP) (*2, 3*)) or by proteasomal degradation of mature proteins (*1, 4*). Antigenic 8-10 amino acid peptides are translocated into the endoplasmic reticulum by the transporter-associated proteins for loading onto MHC I, which are then directed to the plasma membrane for presentation and T cell recognition (*1, 5, 6*). Cell-mediated immunity (CMI) responses generated by this mechanism are critical to preventing disease or reducing severity for respiratory influenza, COVID-19 and respiratory syncytial viral diseases (*7–9*).

Persistent viruses are under selection pressure to evade T-cell responses (*10, 11*). One way that viruses achieve this is by evolving primary amino acid sequences that reduce MHC I immunologic proteolytic processing, which we refer to as protein-intrinsic immune evasion. This generates a cold antigen that does not provoke a CTL priming response. Both Kaposi sarcoma herpesvirus (KSHV) and Epstein-Barr virus (EBV), for example, encode latent proteins with repetitive peptide domains that retard protein translation to maximize correct nascent protein formation and minimize DRiP-mediated antigen presentation (*12–15*). These two viral proteins are also remarkably stable which limits proteasomal processing of the mature proteins (*16, 17*). For AIDS-KS patients, failure to generate effective T-cell responses against latent KSHV proteins can lead to persistent and refractory KS even after antiretroviral therapy has induced immune reconstitution (*18, 19*).

Current RNA-based vaccines for severe acute respiratory syndrome coronavirus 2 (SARS-CoV-2) rely on generating neutralizing antibodies to spike (S) entry protein (*20–23*). Viral escape variants of concern (VoC) with mutations at key neutralizing epitopes (*24*) markedly reduce vaccine efficacy and have led to additional waves of SARS-CoV-2 infection and disease among previously protected vaccinees (*25–27*). While the original BNT162b2 vaccine was 96% effective in preventing disease by the alpha variant, vaccine efficacy dropped to 87% against the delta variant and 31% against the omicron variant (*28*). At this juncture, effective COVID-19 vaccine prevention using S protein vaccines requires reformulation and revaccination once a new, widely circulating VoC is identified.

SARS-CoV-2, an RNA virus, is highly prone to sequence variation throughout most of its genome. Development of a universal SARS-CoV-2 vaccine effective against a broad range of VoC can only be achieved by targeting the small number of conserved non-structural proteins that are essential for virus replication and have particularly constrained mutability. The RNA-dependent RNA polymerase (RdRp or Nsp12) is such a protein, which is conserved among SARS-CoV-2 VoC strains with >99% amino acid identity (**Fig. 1A**) (*29–31*). This essential replication protein does not have a human homolog and is also highly conserved with RdRp from other human betacoronaviruses (SARS-CoV, SARS-CoV-2 and Middle East respiratory syndrome coronavirus, MERS-CoV) rendering it an appealing vaccine target to prevent emergence of new betacoronavirus infections in human populations as well (**Fig. 1B, S1**) (*32, 33*). RdRp, however, is an intracellular protein that is not susceptible to antibody neutralization and can only be targeted by T-cell based immunization.

**Fig. 1.**
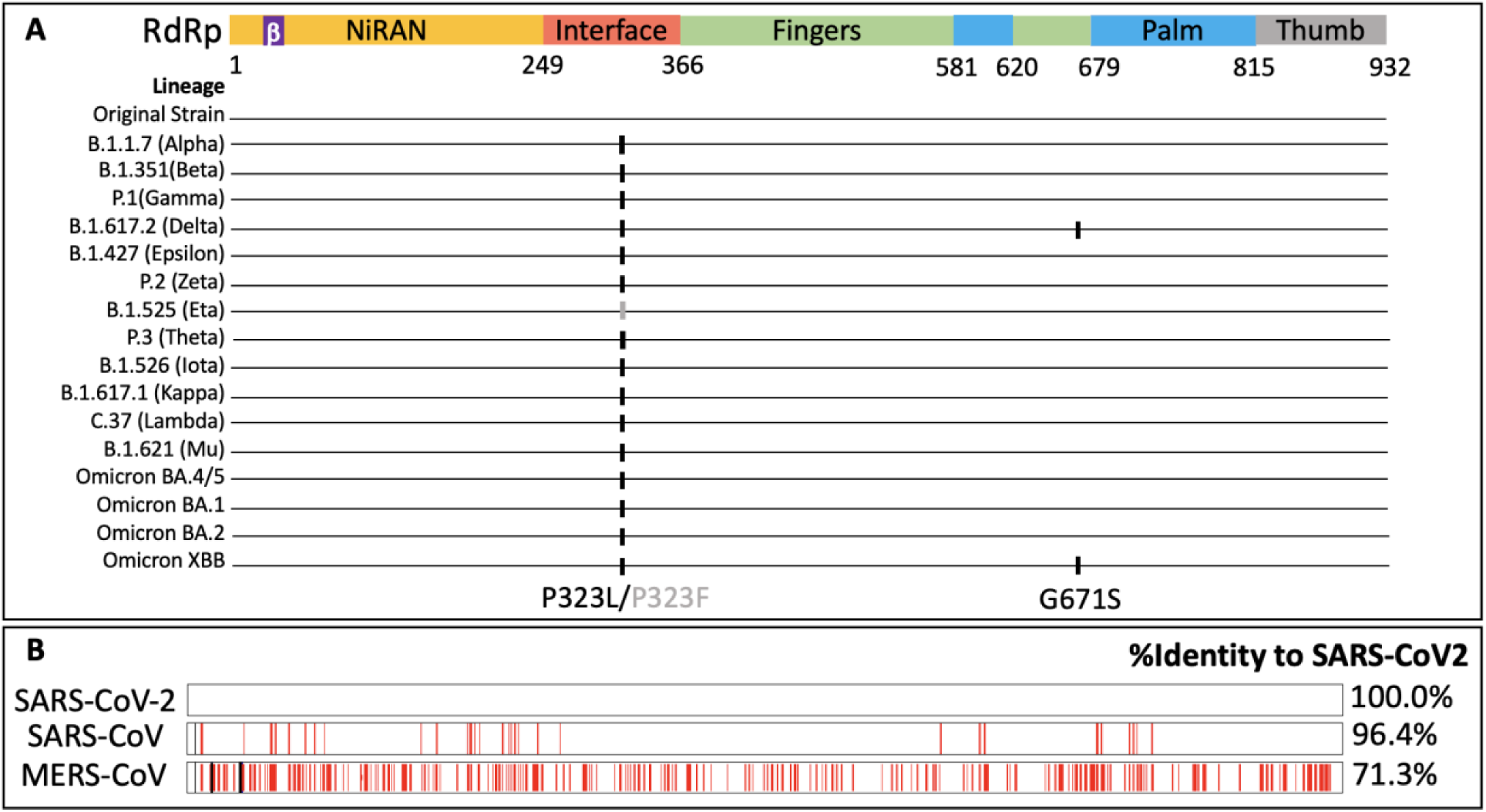
RdRp is conserved among SARS-CoV-2 strain variants and among other human betacoronaviruses. **(A) Amino acid substitutions of the SARS-CoV-2 RdRp protein in SARS CoV-2 variants of concern (VoC).** The primary structure of RdRp with protein domains shown in color: NiRAN (nucleotidyltransferase) at the N-terminus (orange) flanking a *β*-like hairpin structure (purple); Interface (red); fingers (green); palm (blue); and thumb (grey). Only two amino acid polymorphisms are documented in the RdRp protein of VoC through to Omicron XBB. All variants have P323L except for B.1.525 (Eta) which harbors a P323F substitution. Two variants, B.1.617.2 (Delta) and omicron XBB, have G671S substitutions. Data were collected from Outbreak.info and Stanford database websites (https://covdb.stanford.edu/variants/omicron_ba_1_3/). **(B) Alignment of human betacoronaviruses SARS-CoV-2, SARS-CoV and MERS-CoV RdRp proteins.** Red bars indicate mismatch sequence and black bars indicate insertion sequence, relative to the SARS-CoV-2 RdRp. SARS-CoV RdRp and MERS-CoV RdRp show 96% and 71% amino acid identity, respectively to SARS-CoV-2 RdRp. Graph was generated by NCBI Align Sequences Protein BLAST (*66*).

Natural SARS-CoV-2 infection can induce a CD8^+^ T cell response against RdRp and Nsp13 to reduce subsequent endemic coronavirus infections (*34*); a response that is not seen by COVID-19 vaccination against spike proteins. Cross-protective coronaviral T cell responses against SARS-CoV-2 RdRp from prior cold virus exposure (*35, 36*) also suggest that optimizing CMI immunogenicity against RdRp is a potential strategy to generate a viable universal SARS-CoV-2 vaccine candidate (*37*). Such a vaccine could supplement existing vaccines to promote clearance of persistent virus (*9*), or if, sufficiently effective, might be used alone to achieve protection against SARS-CoV-2 VoC. Recurrent COVID-19 disease, however, demonstrates that natural exposure to wild-type RdRp protein during infection is usually not sufficient to generate robust and lasting protective CTL responses. The disappointing immunogenicity of T cell-based vaccines in humans for other viruses, such as HIV, also raises concerns about how the T cell arm of the immune system can be best primed to elicit protection (*38*). Artificially engineering destabilized antigens to enhance peptide presentation is one strategy to prime an initial CTL response that may prove protective during subsequent natural exposure. While licensed mRNA-based COVID-19 vaccines were originally designed to provoke neutralizing S antibodies (*39, 40*), mRNA-delivered antigens effectively generate CD4^+^ processed antigen presentation and holds even greater promise to generate protective T-cell based immunity if a sufficiently antigenic candidate protein sequence can be identified.

How, then, can cold viral proteins be turned into hot antigens while still preserving primary peptide sequences needed for an effective immune response? In this study, we demonstrate the efficacy of protein engineering to overcome viral protein-intrinsic immune evasion to enhance CD8^+^ T cell epitope priming. To show that this can be a general approach to CTL vaccine design, we generated vaccine candidates against conserved replication proteins from an RNA respiratory virus (SARS-CoV-2 RdRp) and a DNA tumor virus (KSHV LANA). We found that the wild-type coding sequence of SARS-CoV-2 RdRp protein, like KSHV LANA, has intrinsic T-cell immune evasion properties suppressing MHC peptide presentation. This can be overcome by dividing the protein into fragments (RdRp^Frag^) to disrupt the stability of nascent protein folding. RdRp^Frag^ had superior T cell immunogenicity over conjugating native RdRp to either ubiquitin or PEST degradation sequences (*41, 42*). For KSHV LANA (**Fig. S2**), deletion of a previously-described inhibitory central repeat (CR) 1 domain, LANA^ΔCR1^, similarly enhanced antigen peptide presentation compared to the native wild-type protein (*43*). Even relatively minor increases (∼2.4 fold) in engineered viral protein MHC I processing and presentation were sufficient to restore specific cytotoxic T cell targeting and tumor growth inhibition for cancer cells stably expressing these vaccine candidates in a syngeneic mouse cancer model. T cells from mice immunized with either the LANA or the RdRp vaccine candidate generated specific CD8^+^ T cell responses on ELISpot assays to target cells expressing the corresponding wild-type proteins. This suggests that CD8^+^ T cell priming by modified LANA and RdRp vaccine candidate antigens can uncover specific viral T cell epitopes that might be recognized during virus infection, despite processing evasion by the native viral protein.

## RESULTS

### *In vitro* KSHV LANA and SARS-CoV-2 RdRp CD8^+^ epitope presentation

To simultaneously determine protein expression level and MHC class I presentation of viral antigens, we generated antigen-EGFP_OVA_ fusion cassettes based on the approach of Yewdell et al. (*3*) (**Fig. 2A**). This produces a single fusion protein containing the virus antigen together with EGFP and the SIINFEKL peptide derived from chicken ovalbumin (OVA), a murine MHC I H2-K^b^-restricted CD8^+^ epitope. This construct design allows simultaneous measurement of protein expression by EGFP fluorescence and H2-K^b^-restricted MHC I SIINFEKL presentation on the cell surface using the 25-D1.16 antibody (*43*). These constructs were then expressed in human 293 cells stably expressing the murine H2-K^b^ MHC I receptor (293KbC2 cells). 293KbC2 cells have a high transfection efficiency and use human processing machinery to present OVA to murine MHC I (*44*). Expression from these constructs was confirmed by immunoblotting (**Figs. S3A, S3B**). To evaluate the OVA presentation in this transfection system, we define the presentation ratio (PR) as the proportion of OVA+ cells within the population of GFP+ cells (**Fig. 2B**). As previously described (*43*), LANA inhibited MHC I OVA peptide presentation (PR = 1.84%) and this was partially reversed by deleting the CR1 region (PR = 4.42%, **Fig. 2C**).

**Fig. 2.**
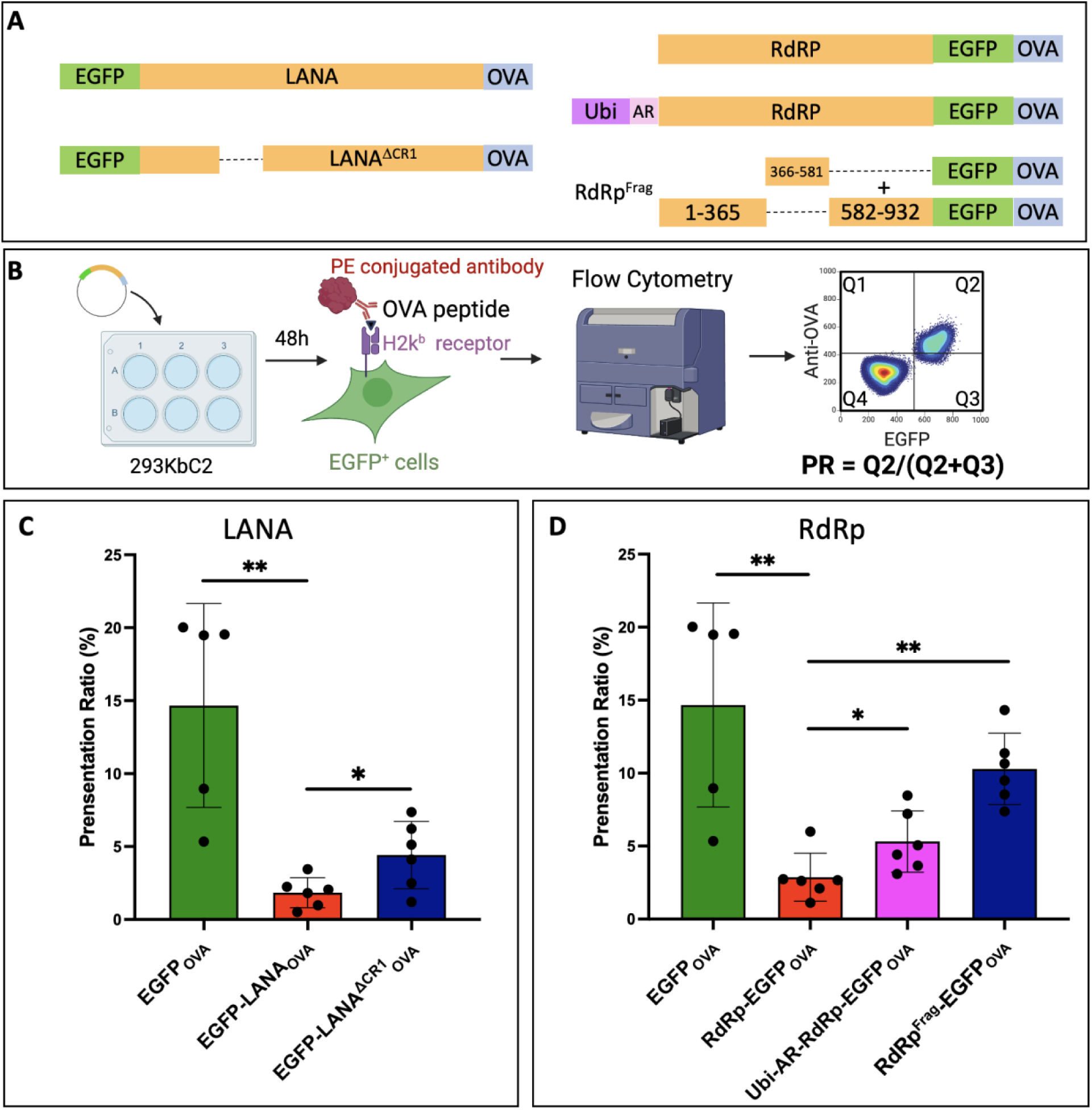
Modified LANA and RdRp promote their MHC I peptide presentation. **(A) Protein engineering of LANA and RdRp constructs used in peptide presentation experiments.** Each viral antigen was fused to EGFP and the chicken ovalbumin peptide SIINFEKL (OVA). **(B)** Wild-type and engineered antigens were transfected into 293 KbC2 cell line, incubated for 48h, and immunostained with PE-conjugated H2-K^b^-OVA specific antibody for flow cytometry. The presentation ratio (PR) was defined as the fraction of cells presenting SIINFEKL over the total GFP^+^ cells (Q2/(Q2+Q3)) to compare MHC class I peptide presentation efficiency. **(C) Increased PR for CR-1 deleted LANA antigens.** EGFP-LANA_OVA_ (red bar) shows significantly decreased PR values compared to EGFP_OVA_ (green bar) consistent with intrinsic CTL immune evasion. This decrease is partially reversed for EGFP-LANA^ΔCR1^_OVA_ (blue bar). **(D) Increased PR for ubiquitinated and fragmentated RdRp antigens.** PR values for native RdRp-EGFP_OVA_ are also reduced (red bar) and can be partially restored by Ubi-AR-RdRp-EGFP_OVA,_ (purple bar) and even greater peptide presentation for RdRp^Frag^-EGFP_OVA_ (blue bar). * P < 0.05, ** P < 0.01, *** P < 0.001 and **** P < 0.0001, Student T test.

Codon-optimized wild-type SARS-CoV-2 RdRp fused to EGFP_OVA_ (RdRp-EGFP_OVA_) also showed diminished OVA presentation (PR = 2.87%) compared to EGFP_OVA_ (PR = 14.67%, **Fig. 2D, S3B**). This is consistent with the notion that the RdRp protein sequence, similar to LANA, has protein-intrinsic immune evasion properties. The estimated half-life of RdRp protein is only 4.2 h on cycloheximide immunoblotting (**Fig. S3C, S3D)** suggesting that RdRp peptide presentation resistance is most likely due to inhibition of nascent rather than mature protein processing.

### Engineering destabilized RdRp promotes MHC I peptide presentation

To increase RdRp destabilization, ubiquitin was directly fused at the N-terminus (*45–47*) with a deubiquitination enzyme (DUB) cleavage site to expose an arginine for proteasome processing through the N-end rule (Ubi-AR-RdRp-EGFP_OVA_) (*48–51*) (**Fig. 2A, S4A**). This increased OVA presentation 1.9-fold over unmodified wild-type RdRp protein (**Fig. 4D**) (N-terminal ubiquitin fusion without the DUB cleavage site resulted in a modest increase in OVA presentation (**Fig. S4A**) in 293KbC2 cells). Cloning a PEST sequence into the C-terminus, either before or after the EGFP fusion peptide, reduced RdRp half-life 3-fold (**Fig. S3D**) but only modestly increased OVA presentation (1.8 to 2-fold) (**Fig. S4B**).

Deletions of specific RdRp regions did not notably increase peptide presentation (**Fig. S5**) suggesting that, unlike LANA (*43*), RdRp does not possess a single discrete peptide domain responsible for CTL immune evasion (**Fig. S5C**). Since mature RdRp protein is not notably stable, we considered it likely that newly-translated RdRp protein folds into a stable structure to avoid DRiP formation.

A strategy was devised to enhance nascent RdRp protein misfolding by sub-fragmenting the protein (*52–54*). A total of twelve different RdRp deletion fragments were generated (**Fig. S5A**) with each fragment cloned in fusion to EGFP_OVA_. Most fragments showed increased OVA presentation compared to full-length wild-type protein with shorter fragments generally displaying higher levels of OVA presentation (**Fig. S5A-C**). We identified a combination of two constructs having the highest levels of OVA presentation normalized by total protein expression (**Fig. S5D**). This combination, called RdRp^Frag^, is comprised of one construct with a deleted “finger” domain (Δ366-581aa) and a second construct encoding the deleted sequence. When these two constructs are expressed in trans, all potential RdRp epitopes are present except those spanning deletion boundaries. Expression of these two constructs together in 293KbC2 cells yielded a 3.6-fold increased OVA presentation compared to the parental wild-type RdRp protein (1-932aa) (**Fig. 2D**).

Although total expression was low in H2-Kb-positive B16-F10 melanoma murine cells, enhanced peptide presentation was confirmed, which showed 9.9-fold higher OVA presentation for the RdRp^Frag^-EGFP_OVA_ combination compared to wild-type RdRp-EGFP_OVA_ (**Fig. S6**). Enhanced OVA MHC I presentation for both LANA^ΔCR1^ and RdRp^Frag^ was also confirmed by measuring antigen presentation over a range of DNA transfection doses in 293KbC2 cells (**Fig. S7A, S7B**). Peptide presentation increased linearly at all levels of LANA^ΔCR1^ and RdRp^Frag^ expression compared to the corresponding wild-type viral proteins.

### Augmented T-cell killing of viral antigen-expressing cells by activated OT-1 CD8^+^ cells

To determine if increased viral antigen presentation results in CD8^+^ T cell cytotoxicity, we used activated OT-1 effector CD8^+^ T cells targeting the OVA peptide epitope (**Fig. 3A**). OT-1 CD8^+^ cells are derived from the OVA-recognizing T cell receptor (TCR) C57BL/6 transgenic OT-1 mouse and were activated by 250ng/ml OVA peptide (Anaspec) for 24 h (*55*). The EGFP_OVA_ constructs were lentivirally transduced into murine colon adenocarcinoma H2-K^b^-restricted MC38 cells and stably selected for similar levels of antigen expression by EGFP cell sorting (**Fig. S8**). Cells were plated onto xCELLigence RTCA DP plates and allowed to adhere for 5 h prior to addition of activated OT-1 CD8^+^ effector cells at various effector: target (E:T) cell ratios.

**Fig. 3.**
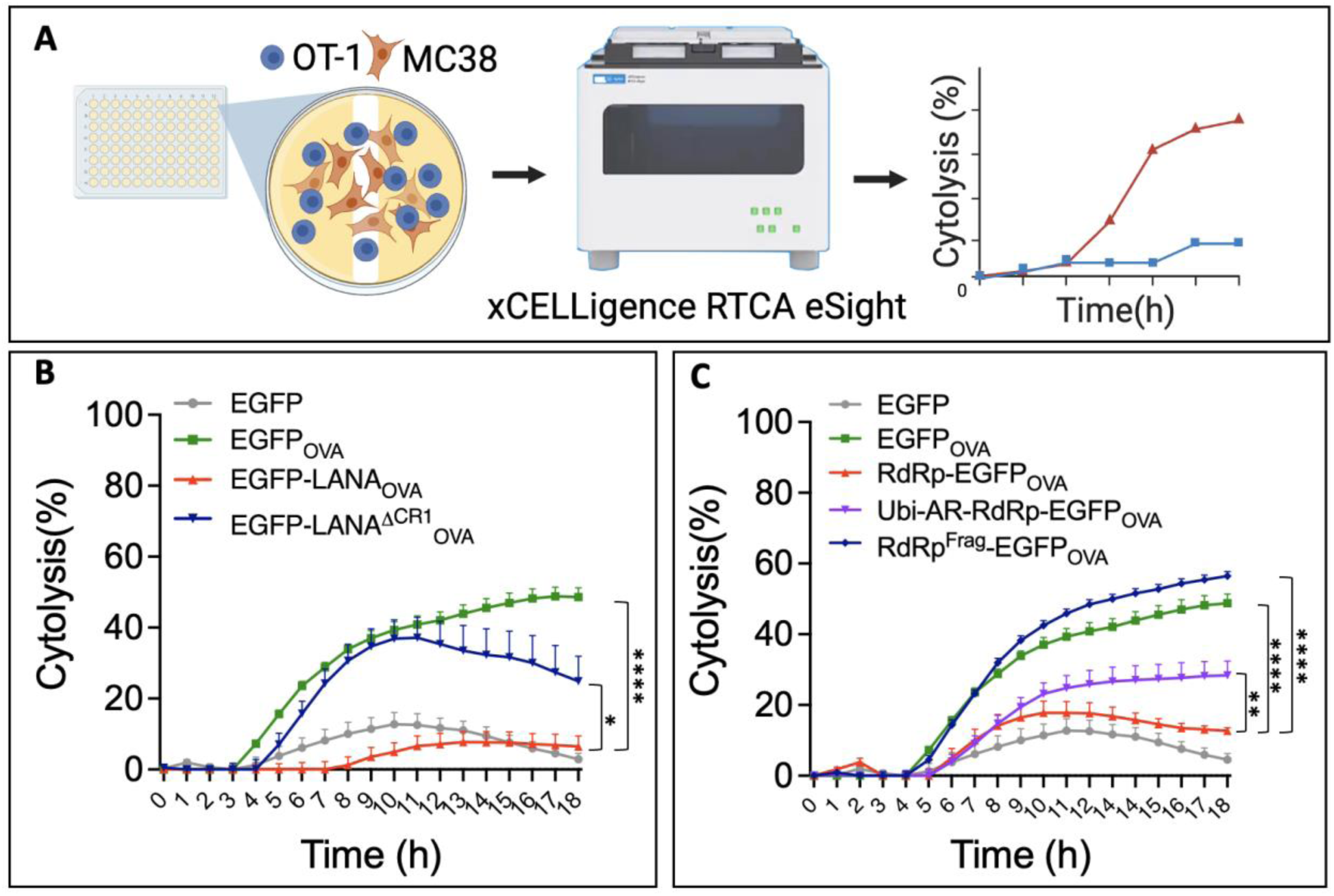
Modified LANA and RdRp enhance CD8+ T cell killing on OT-1 cytotoxic T lymphocyte (CTL) assays. **(A) CTL co-culture assay.** OT-1 cells (CD8*^+^* T cells specifically targeting OVA peptide) were co-cultured with MC38 colon cancer cells stably expressing target antigens fused to OVA on gold-coated 96 well plates. XCELLigence RTCA eSight was used to monitor the impedance for each well in real time. Results were converted into percent cell cytolysis, reflecting OT-1 cell killing efficiency. **(B) Increased cytolysis of MC38 cells expressing LANA^ΔCR1^ by OT-1 cells.** EGFP-LANA_OVA_ showed reduced OT-1 cytolysis (red line) comparable to the negative control cells (grey line, EGFP alone) consistent with intrinsic immune evasion. Expression of EGFP-LANA^ΔCR1^_OVA_ (blue line) restored cytolysis to levels similar to positive control cells (green line, EGFP_OVA_). Effector (OT-1): target cell (MC38) plating ratio = 1:10. A two-way ANOVA test shows a significant interaction between the effect of antigen and time (F(54,209)=8.767, p<0.0001), and a significant effect of antigen on cytolysis (p<0.0001). * P < 0.05, ** P < 0.01, *** P < 0.001 and **** P < 0.0001 determined with post-hoc Tukey test at 18h. N=4 per group. **(C) Increased OT-1 cytolysis of MC38 cells expressing ubiquitin fusion and RdRp^Frag^ antigens.** RdRp^Frag^-EGFP_OVA_ expressing MC38 cells induce the highest target CTL response (blue line) compared to positive control (green line, EGFP_OVA_) and Ubi-AR-RdRp-EGFP_OVA_ cells (purple line). Cytolysis for native RdRp-expressing cells (red line) was significantly greater than the negative control cell line (grey line, EGFP alone). Effector (OT-1): target cell (MC38) ratio = 1:10. A two-way ANOVA test shows a significant interaction between the effect of antigen and time (F(72,285)=14.52, p<0.0001), and a significant effect of antigen on cytolysis (p<0.0001). * P < 0.05, ** P < 0.01, *** P < 0.001 and **** P < 0.0001 determined with post-hoc Tukey test at 18h. N=4 per group. Each data point represents mean± SD.

#### LANA cytotoxicity

As seen in Fig. 3B, EGFP_OVA_ positive control cells (orange line) showed prompt, early OT-1-induced cytotoxicity that was sustained throughout the 18 h time course at E:T=1:10. Low level, nonspecific cytotoxicity occurred for negative control cells, starting at 8 h incubation (EFGP alone, light blue line). Expression of KSHV LANA (EGFP-LANA_OVA_, red line) showed prompt but reduced cytotoxicity compared to EGFP_OVA_, consistent with CTL-immune evasion by this protein. LANA cytotoxicity did not significantly differ from the negative control cells. In contrast, MC38 cells expressing EGFP-LANA^ΔCR1^_OVA_ (dark blue line) had progressive cell killing comparable to the EGFP_OVA_ positive control which remained significantly greater than that of EGFP-LANA_OVA_ by 6 h. These results were consistent over different E:T ratios (**Fig. S9**).

The same assay was performed on 293KbC2 cells transiently transfected with the LANA antigens for cross-validation in a human cell line. Fig. S9 shows a similar trend for 293KbC2 cell cytolysis, where EGFP-LANA^ΔCR1^_OVA_ (navy line) significantly enhanced cytolysis compared to EGFP-LANA_OVA_ (salmon line) at ET= 1:50 (**Fig. S10**).

#### RdRp cytotoxicity

Similar results were obtained for SARS-CoV-2 RdRp at E:T =1:10, with greatest cell killing occurring for RdRp^Frag^ expressing cells (dark blue line, **Fig. 3C**). Wild-type RdRp (red line) showed suppressed cytotoxicity similar to or slightly lower than that of the negative control EGFP alone lacking SIINFEKL epitope (green line), consistent with intrinsic CTL immune evasion for the native RdRp. The ubiquitin-fused RdRp antigen (Ubi-AR-RdRp-EGFP_OVA_, green line) showed modestly increased cytotoxicity that reached significance by 12 h. RdRp^Frag^ expressing cells (dark blue line) revealed the highest CD8^+^ recognition and cell killing throughout the time course, significantly exceeding the cytotoxicity for positive control cells (EGFP_OVA_, green line). Both the Ubi-RdRp fusion protein and the RdRp^Frag^ had significantly increased cell killing at all E:T ratios tested, with RdRp^Frag^ consistently demonstrating a greater effect than Ubi-RdRp (**Fig. 3C, S11**).

### CD8**^+^** T cell responses against either LANA or RdRp suppress syngeneic tumor growth

SARS-CoV-2 is not known to be associated with human cancer but tumor models in syngeneic mice allow us to measure *in vivo* CD8^+^ T cell responses to processed antigens. To measure T cell responses in mice, naïve 8-week-old C57BL/6 mice were subcutaneously injected in the flank with 2.5×10^5^ syngeneic MC38 cells expressing control (EGFP_OVA_), wild-type or modified LANA/RdRp proteins. Tumor cells expressing native LANA-EGFP_OVA_ and RdRp-EGFP_OVA_ proteins generated tumors that required sacrifice starting at ∼12-13 days whereas survival was significantly extended (>20 days) for mice receiving candidate vaccine antigens or the EGFP_OVA_ cells.

Figures **4A** and **4B** show survival and tumor growth curves for EGFP_OVA,_ EGFP-LANA_OVA_ and EGFP-LANA^ΔCR1^_OVA_ groups. Mice injected with tumor cells expressing the EGFP_OVA_ or EGFP-LANA_OVA_ developed palpable tumors that regressed by day 7. Tumors reemerged in both EGFP_OVA_ and EGFP-LANA_OVA_ groups after day >15, which resulted both groups having a median survival of >34 days when the experiment was ended (**Fig. 4B**, green line).

**Fig. 4.**
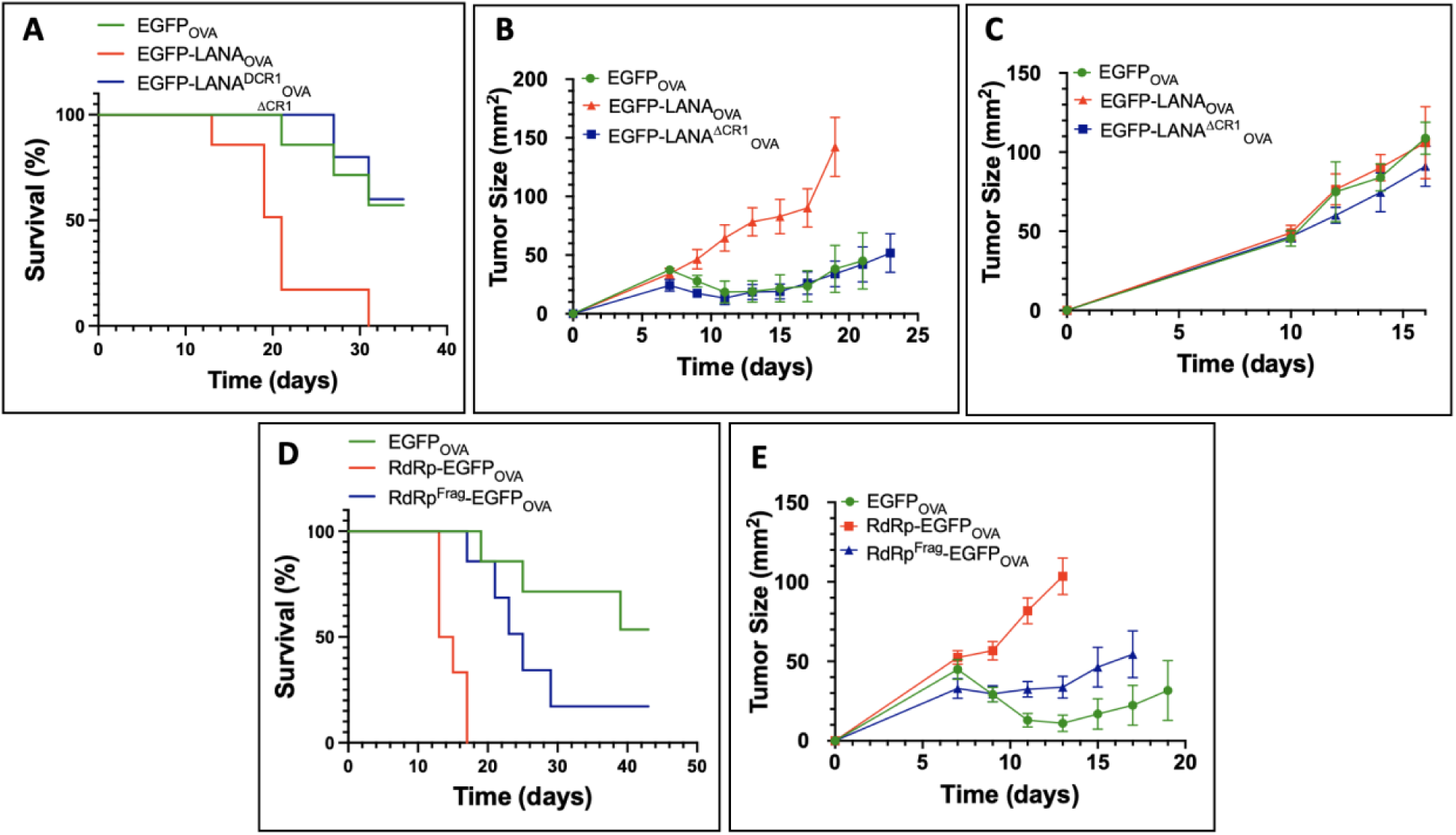
Survival and tumor growth rates for mice injected with MC38 cells expressing LANA and RdRp antigens. **(A) Kaplan-Meier survival curve for C57BL/6 mice injected with MC38 cells expressing LANA antigens.** EGFP-LANA_OVA_ causes early mortality (red line), with a median survival of 19 days. EGFP-LANA^ΔCR1^_OVA_ (blue line) shows delayed mortality (median 34 d) that is comparable to EGFP_OVA_. Log-rank test for EGFP-LANA_OVA_ vs. EGFP-LANA^ΔCR1^_OVA_, P= 0.0046. N=6 per group. **(B) Tumor size growth curve for C57BL/6 mice injected with MC38 cells expressing LANA antigens.** Expression of EGFP-LANA_OVA_ in MC38 colon cancer cells results in more rapid tumor growth compared to LANA^ΔCR1^_OVA_ or EGFP_OVA_. Two-way ANOVA test for data from day 0 to day 19 shows a significant interaction between the effect of antigen and time (F (14,98)= 8.458, p<0.0001), and a significant effect of antigen on tumor size (p=0.0036). N=6 per group. **(C) Tumor size growth in Rag1 KO mice injected with MC38 cells expressing various LANA antigens.** Rag1 KO (B6.129S7-Rag1^tm1Mom^/J) mice were monitored for 16 days. No differences in tumor growth were detected between different groups, two-way ANOVA, p=0.1321. N=5 per group. **(D) Kaplan-Meier survival curve for C57BL/6 mice injected with MC38 cells expressing RdRp antigens.** MC38 cells expressing RdRp-EGFP_OVA_ (median survival 13 D, red line) have significantly greater early mortality compared to cells expressing RdRp^Frag^-EGFP_OVA_ (median survival 23 d, blue line), log-rank test P=0.001, n=6 per group. **(E) Tumor size growth curve for C57BL/6 mice injected with MC38 cells expressing RdRp antigens.** A two-way ANOVA test for data from day 0 to day 13 shows a significant interaction between the effect of antigen and time (F(8, 68)= 25.27, p<0.0001), and a significant effect of antigen on tumor size (p<0.0001), n=6 per group. Each data point represents the Mean ± SD.

In contrast to the EGFP_OVA_ group, mice injected with EGFP-LANA_OVA_ cells showed no tumor regression (**Fig. 4B**, red line) and reached median survival by day 20, consistent with LANA-SIINFEKL protein evasion of CTL recognition. Tumor growth and survival for mice injected with EGFP-LANA^ΔCR1^_OVA_ cells (**Fig. 4B**, blue line), however, were similar to those of the EGFP_OVA_ control group, and showed significantly increased survival compared to the EGFP-LANA_OVA_ group (log-rank test, EGFP-LANA_OVA_ vs. EGFP-LANA^ΔCR1^_OVA_ p=0.0046). To demonstrate that the antitumoral activity of LANA^ΔCR1^ is related to CD8^+^ T cell immunity, this experiment was repeated in C57BL/6 Rag1 knockout mice lacking effective CD8^+^ T cells. All three antigen-bearing MC38 tumors grew indistinguishably up to the experimental endpoint (**Fig. 4C**).

Mice injected with RdRp antigen-MC38 cells showed similar patterns of survival and tumor formation (**Fig. 4D and 4E**). Monotonic increase in tumor size and reduced survival were noted for mice in the RdRp-EGFP_OVA_ group compared to RdRp^Frag^-EGFP_OVA_ and EGFP_OVA_ groups with mice in the RdRp group having a median survival of 13 days compared to 23 days for those in the RdRp^Frag^ group (log-rank test, RdRp-EGFP_OVA_ vs. RdRp^Frag^-EGFP_OVA_, p=0.001). Tumor growth curves for MC38 tumors expressing RdRp^Frag^-EGFP_OVA_ (blue line) were intermediate between RdRp-EGFP_OVA_ (red line) and EGFP_OVA_ (green line).

To determine if the vaccine candidate proteins elicited T cell recognition of viral epitopes in addition to the SIINFEKL epitope, ELISpot assays were performed using MC38 target cells expressing the native viral proteins without the EGFP_OVA_ fusion peptide (**Fig. S12**). **Fig. 5** shows γ-interferon (IFN-γ) ELISpot counts for CD8^+^ T cells harvested from single surviving LANA^ΔCR1^-, RdRp^Frag^- and OVA-injected mice (day 90 after initial injection) coincubated with the MC38 target cells. Both wild-type LANA and RdRp proteins were capable of activating CD8^+^ T cells from mice exposed to their corresponding vaccine candidate (**Fig. 5B**).

**Fig. 5.**
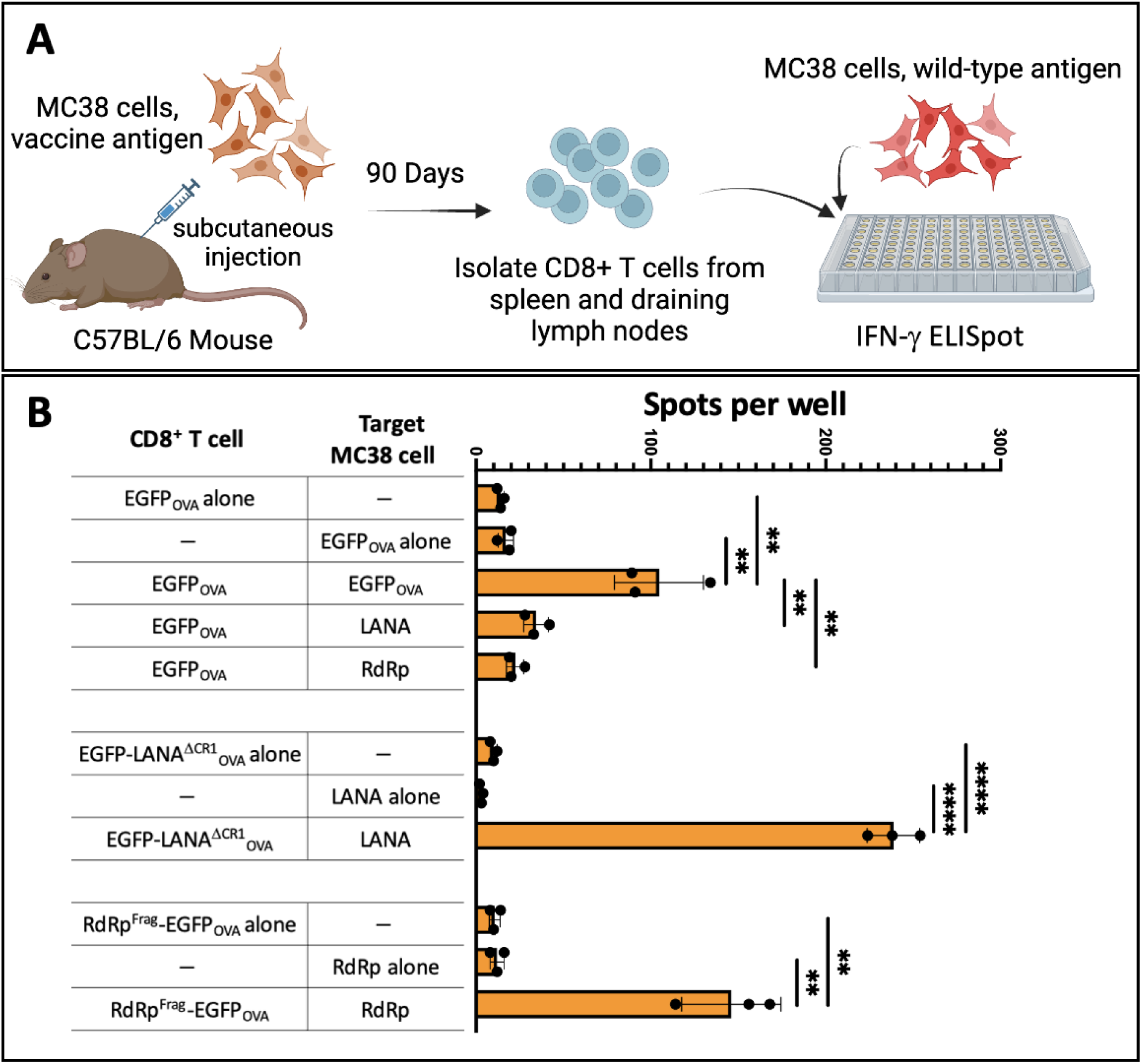
Modified LANA and RdRp immunized mice generate immune memory to viral antigens in CD8^+^ T cells. **(A) Experimental Setup for ELISpot assay.** Individual C57BL/6 mice were subcutaneously injected with MC38 cells expressing EGFP_OVA,_ EGFP-LANA^ΔCR1^_OVA_ or RdRp^Frag^-EGFP_OVA_ to form tumors that subsequently regressed. Rare surviving mice were harvested 90 days post injection for the collection of CD8^+^ T cells from draining lymph nodes and spleen. CD8^+^ T cells were then co-cultured with MC38 cells expressing wild-type LANA or RdRp proteins lacking the OVA tag to assess immunogenicity using an ELISpot assay. **(B) LANA^ΔCR1^_OVA_ and RdRp^Frag^_OVA_ immunized mice generated CD8^+^ T cell responses to wild-type viral antigens.** CD8^+^ T cells from a mouse immunized with EGFP_OVA_ (positive control) after 90 days generated T cell IFN-γ secretion only during incubation with target MC38 cells expressing EGFP_OVA_ but not for T cells or MC38 target cells alone or when coincubated with MC38 cells expressing the wild-type viral proteins lacking the EGFP_OVA_ tag. CD8^+^ T cells isolated from mice immunized with EGFP-LANA^ΔCR1^_OVA_ or RdRp^Frag^-EGFP_OVA_ after 90 days, showed T cell activation only when incubated with cells expressing their respective wild-type viral proteins lacking the EGFP_OVA_ tag. Error bars indicate mean ± SD from triplicate wells. * P < 0.05, ** P < 0.01, *** P < 0.001 and **** P < 0.0001 as determined by Student T test.

### CD8^+^ T cell responses to modified LANA and RdRp proteins in mice

To measure CD8^+^ T cell responses to antigen presentation, a cohort of mice were harvested eight days after injection to obtain tumor-infiltrating lymphocytes (TIL) as well as CD8^+^ T cells from draining lymph nodes and splenic tissues. These cells were then stained and examined using the OVA TCR tetramer (PE-H2-K^b^/OVA (SIINFEKL) MHC Tetramer, 1:100, NIH Tetramer Core Facility) by flow cytometry (**Fig. S13**).

#### LANA CD8^+^ T cell responses

EGFP-LANA_OVA_ cell injection suppressed total OVA-specific CD8^+^ T cell staining in all tissue compartments compared to EGFP_OVA_ alone (**Fig. 6A**). Activated effector T cells (CD44^+^/CD62L^-^) similarly were suppressed by EGFP-LANA_OVA_ cell injection (**Fig. 6B**). Although total CD8^+^ T cell suppression was significantly, but modestly, reversed in the lymph node and spleen tissues for mice receiving EGFP-LANA^ΔCR1^_OVA_ cell injections, T effector cells (CD8^+^/CD44^+^/CD62L^-^) showed a marked reversal of TIL suppression (**Fig. 6A**). Specific T effector cells were not statistically different in spleen tissues for mice injected with LANA or LANA^ΔCR1^ cells at this early time point.

**Fig. 6.**
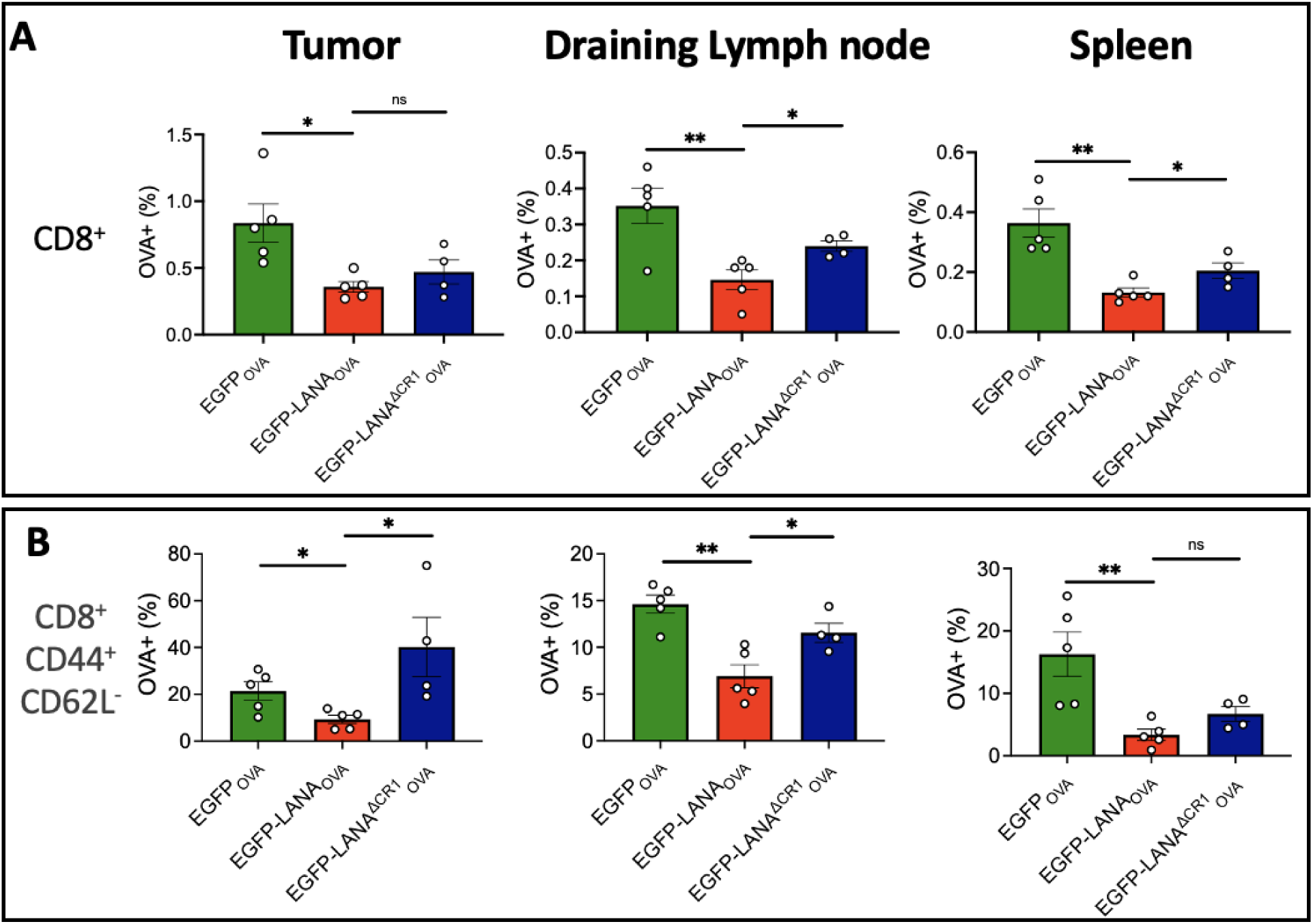
Increased OVA specific T cell responses in mice injected with MC38 cells expressing EGFP-LANA^ΔCR1^_OVA_ from tumor (left column), draining lymph node (middle column), and spleen (right column). **(A) Percentage of OVA specific T cells among CD8^+^ T cell populations** show EGFP-LANA_OVA_ repressed OVA specific T cell response and effective recovery by EGFP-LANA^ΔCR1^_OVA_ for draining lymph node and spleen, but not for tumor tissue. **(B) Percentage of OVA specific effector T cells among CD8^+^ effector T cells (CD44^+^/CD62L^-^)** shows EGFP-LANA_OVA_ repressing OVA specific effector T cell response and effective recovery by EGFP-LANA^ΔCR1^_OVA_ for tumor and draining lymph node, but not for spleen. * P < 0.05, ** P < 0.01, *** P < 0.001 and **** P < 0.0001 as determined by Student T test (n=5 per group).

#### RdRp CD8^+^ T cell responses

For the native RdRp protein group, like the native LANA protein group, OVA tetramer staining was suppressed for TIL, draining lymph nodes and spleen by day 8 (**Fig. 7A**). This was unchanged after Ubi-RdRp-EGFP_OVA_ cell injection. RdRp^Frag^-expressing cells markedly increased total CD8^+^ T cell OVA-specific responses in tumor, draining lymph node and spleen tissues. Similar to LANA^ΔCR1^, this immunostimulatory effect was even more pronounced for CD8^+^ effector staining (CD44^+^/CD62L^-^) in all three tissues examined (**Fig. 7B**).

**Fig. 7.**
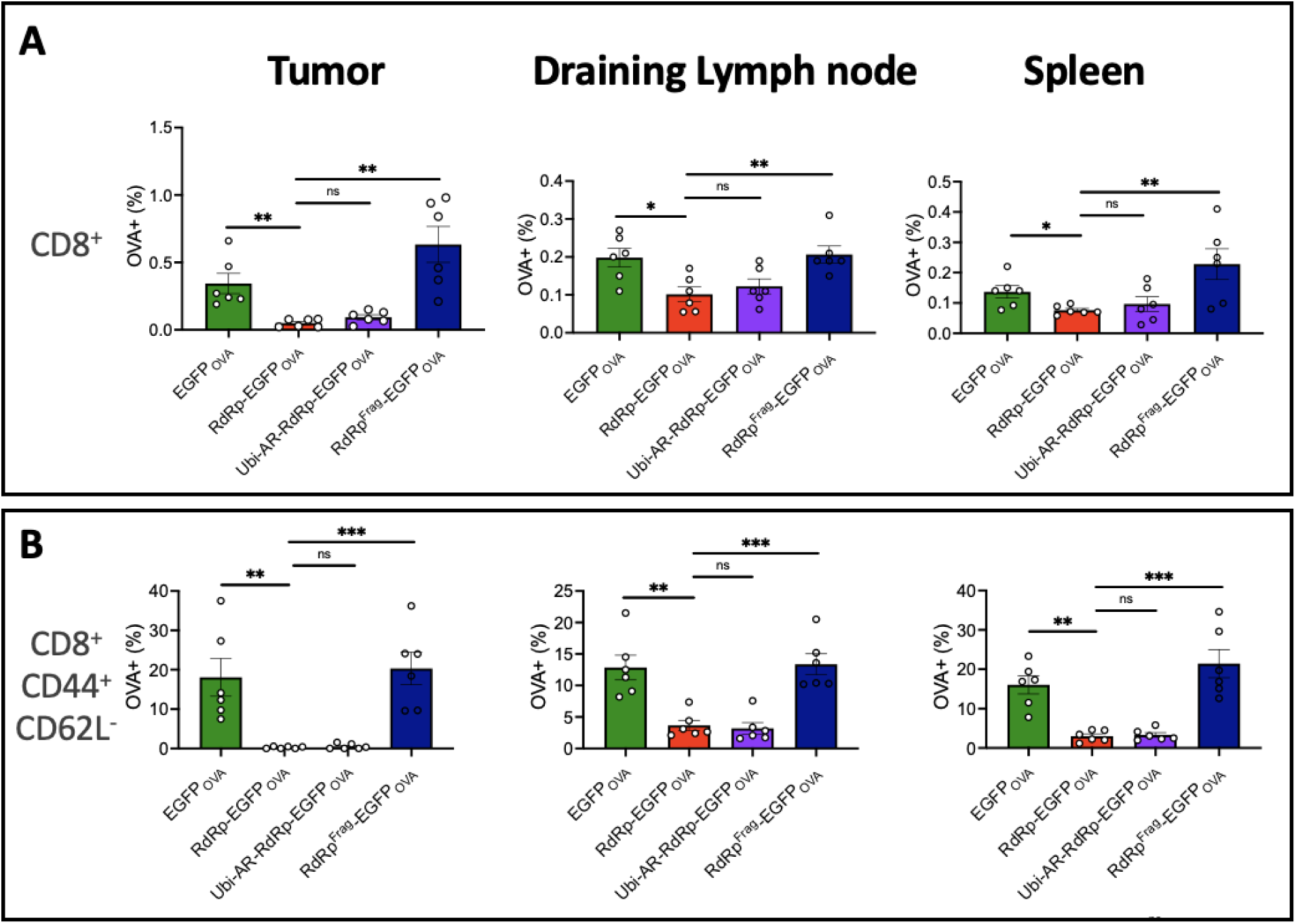
Increased OVA specific T cell response in mice injected with MC38 cells expressing RdRp^Frag^-EGFP_OVA_ from tumor (left column), draining lymph node (middle column), and spleen (right column). **(A) Percentage of OVA specific T cells among CD8^+^ T cell populations** show RdRp-EGFP_OVA_ repressed OVA specific T cell response that is effectively reversed by RdRp^Frag^-EGFP_OVA_ but not by Ubi-AR-RdRp-EGFP_OVA_, for tumor, draining lymph node, and spleen. **(B) Percentage of OVA specific effector T cell percentage among CD8^+^ effector T cells (CD44^+^/CD62L^-^)** shows RdRp-EGFP_OVA_ represses OVA specific effector T cell response that is reversed by RdRp^Frag^-EGFP_OVA_ but not Ubi-AR-RdRp-EGFP_OVA_, for tumor, draining lymph node, and spleen. * P < 0.05, ** P < 0.01, *** P < 0.001 and **** P < 0.0001 as determined by Student T test (n=5 per group).

In summary, injection of MC38 cells expressing either LANA or RdRp suppressed total T cell and T effector cell-induced tetramer staining in TIL and draining lymph nodes. Both, LANA^ΔCR1^ and RdRp^Frag^ proteins reversed this suppression, generating tetramer staining similar to the OVA positive control antigen.

## DISCUSSION

AIDS-KS patients with immune reconstitution after antiretroviral therapy often achieve KS remission, consistent with antigen presentation by tumors (*56*). LANA is an abundant protein in all KSHV-related cancers and LANA strain variations are largely limited to changes in the numbers of central repeats (*57*). If LANA^ΔCR1^ proves useful as a therapeutic vaccine, it would be particularly beneficial for HIV-negative KS patients and those HIV-positive patients having refractory KS after successful antiretroviral therapy. The clinical importance of a CTL vaccine against SARS-CoV-2 RdRp is less certain. While RdRp is essential for SARS-CoV-2 replication, RdRp expression is sparse in lung tissues of patients dying from COVID-19 (*58*). It is also unclear that T cell responses to RdRp alone can either prevent infection or limit severity of disease after infection (*58*). These concerns can be readily addressed by animal challenge experiments. If RdRp^Frag^ has clinical utility in preventing or limiting the clinical severity of COVID-19, it is a vaccine candidate that may be useful against most, if not all, strains of SARS-CoV-2 and closely-related betacoronaviruses.

In this study, we show that two unrelated viral replication proteins, LANA and RdRp, have primary amino acid sequences that evade CD8^+^ T cell processing machinery. Protein intrinsic immune evasion also has been described for EBV EBNA1(*59, 60*), human adenovirus E3/19K(*61*) and CMV US3(*62*). It is likely that similar mechanisms to evade immune processing are more common for viral proteins than generally recognized. Yewdell et al first proposed DRiP degradation of misfolded nascent proteins as a source for CMI peptide presentation (*3, 63*), which is predicted to have a 10-minute half-life after initiation of mRNA translation. *In vitro* presentation assays using a fused OVA (SIINFEKL) tag allowed identification and engineering of antigen sequences with enhanced presentation that retain most viral epitopes needed for CTL responses. We show here that even modest improvement in LANA and RdRp peptide presentation generates potent CTL cell killing and suppression of tumorigenesis in mice.

In the case of SARS-CoV-2, dividing the RdRp protein sequence into two fragments was sufficient to restore immune processing to generate potent CTL responses (**Fig. 2D**). While RdRp is not associated with cancer during natural SARS-CoV-2 infection, the MC38 tumor model is a well-established model for CTL responses in C57BL/6 mice. It is currently unknown whether tumor reemergence after initial regression after immunization results from escape mutations that eliminate the antigens or failure of T cell responses to control MC38 tumor cell outgrowth. ELISpot assays, however, indicate that mice can develop durable T cell recognition to native viral proteins.

Our study supports use of engineered, destabilized viral proteins, and measuring peptide processing *in vitro*, as an approach to improving whole protein CTL immunogenicity by reducing protein-intrinsic T cell immune evasion. The clinical utility of these LANA and RdRp vaccine candidates requires additional preclinical evaluations, but these findings reinforce the value of surrogate antigen presentation assays as a relatively inexpensive and simple approach to screen for optimized whole protein CTL vaccine antigens.

## MATERIALS AND METHODS

### Plasmids

The construction of EGFP_OVA,_ EGFP-LANA_OVA_, EGFP-LANA^ΔCR1^_OVA,_ EGFP-PEST_OVA_ have been previously described (*43*). Codon-optimized RdRp coding sequence (**Table S1**) was synthesized by Integrated DNA Technologies (IDT, Coralville, IA) and cloned as a fusion protein N-terminal to EGFP_OVA_ and EGFP-PEST_OVA_ constructs at *BamH*I/*EcoR*I sites to generate RdRp-EGFP_OVA_ and RdRp-EGFP-PEST_OVA_, respectively. RdRp-EGFP_OVA_-PEST was synthesized by GenScript Biotech (Piscataway, NJ). All RdRp fragments were generated by PCR from the synthesized codon-optimized RdRp template and inserted N-terminal to EGFP_OVA_ at *BamH*I/*EcoR*I restriction sites.

Ubi-RdRp-EGFP_OVA_ was generated by PCR of the ubiquitin sequence from pRK-HA-Ubiquitin-WT (Addgene, #17608) which was inserted at the N-terminus of RdRp-EGFP_OVA_ using *Sac*II/*EcoR*I. To generate a modified Ubi-RdRp-EGFP_OVA_ construct incorporating the N-end rule which we named Ubi-AR-RdRp-EGFP_OVA_, a different reverse primer for PCR was used that generated the C-terminus of ubiquitin with G to A mutation immediately followed by an A to R mutation. To generate pMuLE EF.Bla, pENTR EF1 vector was LR-cloned into pMule Lenti DEST Bla, which we constructed from in pMuLE Lenti DEST Neo (Addgene, #62178) by replacing *neo* gene with *bsr* using *Sbf*I and *Xho*I. All plasmids produced for lentiviral transduction were generated by PCR from the respective target gene and inserted into the pLVX EF.Puro plasmid (*64*) at *Afe*I/*Sbf*I sites, except pLenti-RdRp.Δ366-581aa-EGFP_OVA_ which was inserted into the pMuLE EF.Bla plasmid. All plasmids are listed in **Table S2** and were sequence confirmed. All primers are listed in **Table S3.**

#### Cell lines

293KbC2 cells (*43*) stably expressing the mouse class I allele H-2Kb (*44*) were maintained in Dulbecco’s modified Eagle medium (DMEM, ThermoFisher) with 10% FBS supplemented with 0.5 mg/ml G418 (Hyclone). B16-F10 cells (ATCC) and MC38 cells (ATCC) were maintained in DMEM supplemented with 10% FBS. All cell lines were maintained in 37°C and 5% CO2.

To generate MC38 cell lines stably expressing target proteins (e.g. RdRp-EGFP_OVA_), lentiviral transduction was performed using a second generation lentiviral packaging system (*65*). In brief, 293FT cell lines (Invitrogen) were transfected with the target lentiviral plasmid, as well as packaging vectors pMD2.G and pSPax2. Supernatant was harvested 72 h after transfection and used to infect MC38 cells in 6-well plates with addition of polybrene at 6 µg/ml, followed by centrifugation in 2000 rcf for 2 h. MC38 cells were infected for 72 h, then selected in puromycin at 5 µg/ml for stable cell lines.

For the RdRp^Frag^-EGFP_OVA_, dual transduction was performed with a first pLVX EF.puro-RdRp.366-581aa-EGFP_OVA_ transduction selected by puromycin at 5 µg/ml, followed by a pMuLE EF.Bla-RdRp.Δ366-581aa-EGFP_OVA_ transduction selected by blasticidin at 10 µg/ml.

### Transfection and cycloheximide treatment

293KbC2 cells were seeded in 6-well plates and transfected with appropriate sample plasmid combinations to equal 2 µg total DNA and 2 µl of Lipofectamine 2000 (ThermoFisher). At 24 h post-transfection, cells were imaged or collected for immunoblotting or flow cytometry.

For cycloheximide treatment, cycloheximide (CHX, Sigma) was added to a final concentration of 100 µg/ml to inhibit protein synthesis 24 h after transfection. Negative control cells were treated with an equal volume of dimethyl sulfoxide (DMSO) vehicle. All samples were collected 6 h after treatment for immunoblotting.

### Immunoblotting

Total protein was extracted from transfected cells using radioimmunoprecipitation assay (RIPA) lysis buffer (150mM NaCl, 1% NP-40,0.5% DOX,0.1% SDS, 50mM Tris-HCl PH7.4) containing protease inhibitors (0.2 mM Vanadate, 0.3 mM PMSF, 1 mg/mL Leupeptin, 1 mg/mL Pepstatin A, 1 mg/mL Aprotinin). Samples were sonicated with Fisherbrand™ Model 505 Sonic Dismembrator (ThermoFisher) at 20% Amp 4x for 5 seconds each on ice. 2x Laemmli loading buffer (65.8 mM Tris-HCl pH 6.8, 26.3% glycerol, 2.1% SDS, 0.01% bromophenol blue), 10% 2-mercaptoethanol were added to samples which were then separated by SDS-PAGE and transferred to nitrocellulose membranes. Membranes were incubated with overnight at 4°C with primary rabbit polyclonal antibody to EGFP (Santa Cruz, sc-8334) diluted 1:500, followed by incubation with IRD800 conjugated goat anti-rabbit secondary antibody (LI-COR Biotechnology) diluted 1:10,000 and Rhodamine conjugated anti-tubulin antibody (Bio-Rad) diluted 1:10,000 for 1 h at room temperature. A ChemiDoc™ MP Imaging system (Bio-Rad) was used to detect signals. Image J was used for quantitation of immunoblot signals.

### Flow cytometry

293KbC2 cells transfected with target constructs for 48 h in 6-well plates were harvested in 1ml PBS and washed twice with 500 µl PBS. Cells were then stained with phycoerythrin (PE) anti-mouse MHC class I Kb-SIINFEKL (25-D1.16) (eBioscience) diluted 1:30 for 1 h at 4°C, then washed twice with 500 µl PBS and stored in 500 µl PBS for analysis on LSRFortessa cell analyzer (BD Bioscience). For each experiment, gating was performed for positive EGFP fluorescence and positive OVA staining, and 30,000 events were collected and analyzed. All experiments were repeated at least three times for reproducibility, with representative experiments shown.

### xCELLigene real-time killing assay

100,000 MC38 cells stably expressing target antigen were seeded onto the E-Plate 96 and incubated for a minimum of 5 h. Subsequently, primary CD8^+^ T cells, isolated from C57BL/6-Tg(TcraTcrb)1100Mjb/J (Strain #:003831, common name: OT1), were introduced to the E-Plate 96 at various Effector (E) to target (T) ratios. Following this, the system was left undisturbed for a minimum of 17 h to allow the Agilent xCELLigence RTCA DP system (Santa Clara, CA) to record impedance changes. The RTCA Software Pro was utilized for cytolysis calculations.

### Tetramer assay

Subcutaneous injections of 250,000 MC38 cells stably expressing target antigen were administered to C57BL/6 mice. All mice were euthanized on day 7 with tumor sizes ranged from 20 mm² to 40 mm². Tumor, draining lymph nodes, and spleens were surgically isolated from mice. Tumor tissues were further treated with enzymatic digestion using collagenase IV and DNAse. All tissues were then mechanically disrupted through a 70μm cell strainer to obtain single-cell suspensions. All samples were treated with 5% mouse serum for 15 minutes on ice in a blocking step, and then stained with CD8β (BV786, BD bioscience), CD39 (Duha59, Biolegend), CD62L (MEL-14, Biolegend), CD44 (IM7, Biolegend), and Zombie NIR™ for 15 minutes on ice. Following this, samples were washed once in PBS before staining with PE-H2-K^b^-OVA(SIINFEKL)-tetramer (1:100, NIH Tetramer Core Facility) for an additional 15 minutes. All samples were then fixed in 4% Paraformaldehyde. Flow cytometry analysis involved gating for Zombie negative, CD8β, and PE-OVA-tetramer, followed by CD44 and CD62L in CD8β^+^ subpopulation. 100,000 events were collected and analyzed per sample.

### ELISpot assay

Subcutaneous injections of 250,000 MC38 cells stably expressing EGFP_OVA_, EGFP-LANA^ΔCR1^_OVA_, or RdRp^Frag^-EGFP_OVA_ were administered to C57BL/6 mice. Survived mice after 90 days with tumor diminished were euthanized. Draining lymph nodes and spleens were surgically isolated and mechanically disrupted through a 70μm cell strainer into single-cell suspensions and mixed. CD8^+^ T cells were then negatively selected from the mix by incubating on ice for 15 min with biotin conjugated antibodies: CD19 (115504, BioLegend) at 1:1000, CD11c (117304, BioLegend) at 1:1000, Ly-6G/Ly-6C (108404, BioLegend) at 1:1000, CD11b (101204, BioLegend) at 1:1000, TCRγ/δ (118103, BioLegend) at 1:1000, CD45R/B220 (103204, BioLegend) at 1:500, CD49b (108904, BioLegend) at 1:500, CD105 (120404, BioLegend) at 1:500, CD24 (101804, BioLegend) at 1:500, CD16/32(101303, BioLegend) at 1:500, CD25(102004, BioLegend) at 1:1000, and CD4(100508, BioLegend) at 1:500. MojoSort Streptavidin Nanobeads (BioLegend) were then applied for 15 min, followed by magnet pull-down for 5 min to remove unwanted cells.

ELISpot were performed with Mouse IFN-γ ELISpot Kit (EL485, R&D System) according to the manufacturer’s protocol. In brief, the selected CD8^+^ T cells were co-cultured overnight with MC38 cells stably expressing WT LANA, WT RdRp, or EGFP_OVA_ at 500,000 cells/ml each (E:T =1:1), into a 96-well PVDF-backed microplate. The plate was then sequentially treated with biotinylated anti-IFN-γ antibody, alkaline phosphatases conjugated streptavidin, and BCIP/NBT substrate. Developed spots were manually counted for ΙFN-γ producing cells.

## Acknowledgments

We thank Simon Cao, Jinghui Li and Siying Guo for initiating SARS-CoV-2 RdRp cloning and expression, and Steve Reich and Philip Glass for help with the manuscript.

## Author contributions

P.S.M and Y.C. conceptualized this work. L.W and B.X. designed and performed the experiments, analyzed the data, and prepared the figures. M.S contributed to the DNA plasmid cloning. G.D. provided the resource for in vivo mouse experiments. G.D. and M.S. provided key academic input into the study design. P.S.M and Y.C. supervised the experiments. L.W., P.S.M., and Y.C. co-wrote the manuscript. All authors reviewed and edited the manuscript.

## Data and material availability

All data are available in the paper or the Supplementary Materials. All cell lines and plasmids used in this study will be made available to the scientific community by contacting the corresponding author and completion of a material transfer agreement.

## References and Notes

1. J. S. Blum, P. A. Wearsch, P. Cresswell, Pathways of antigen processing. Annual review of immunology 31, 443–473 (2013).

2. U. Schubert et al., Rapid degradation of a large fraction of newly synthesized proteins by proteasomes. Nature 404, 770–774 (2000).

3. J. W. Yewdell, L. C. Antón, J. R. Bennink, Defective ribosomal products (DRiPs): a major source of antigenic peptides for MHC class I molecules? Journal of immunology (Baltimore, Md.: 1950) 157, 1823–1826 (1996).

4. D. Bourdetsky, C. E. Schmelzer, A. Admon, The nature and extent of contributions by defective ribosome products to the HLA peptidome. Proc Natl Acad Sci U S A 111, E1591–1599 (2014).

5. J. Neefjes, M. L. Jongsma, P. Paul, O. Bakke, Towards a systems understanding of MHC class I and MHC class II antigen presentation. Nature reviews immunology 11, 823–836 (2011).

6. K. L. Rock, E. Reits, J. Neefjes, Present yourself! By MHC class I and MHC class II molecules. Trends in immunology 37, 724–737 (2016).

7. C. B. Hall et al., Respiratory syncytial viral infection in children with compromised immune function. New England Journal of Medicine 315, 77–81 (1986).

8. A. Jozwik et al., RSV-specific airway resident memory CD8+ T cells and differential disease severity after experimental human infection. Nature communications 6, 10224 (2015).

9. Y. Li et al., SARS-CoV-2 viral clearance and evolution varies by type and severity of immunodeficiency. Sci Transl Med 16, eadk1599 (2024).

10. T. H. Hansen, M. Bouvier, MHC class I antigen presentation: learning from viral evasion strategies. Nature Reviews Immunology 9, 503–513 (2009).

11. J. L. Petersen, C. R. Morris, J. C. Solheim, Virus evasion of MHC class I molecule presentation. The Journal of immunology 171, 4473–4478 (2003).

12. M. E. Ressing et al., in Seminars in cancer biology. (Elsevier, 2008), vol. 18, pp. 397–408.

13. O. Sorel, B. G. Dewals, The critical role of genome maintenance proteins in immune evasion during gammaherpesvirus latency. Frontiers in Microbiology 9, 3315 (2019).

14. E. W. Hewitt, The MHC class I antigen presentation pathway: strategies for viral immune evasion. Immunology 110, 163–169 (2003).

15. J. Levitskaya, A. Sharipo, A. Leonchiks, A. Ciechanover, M. G. Masucci, Inhibition of ubiquitin/proteasome-dependent protein degradation by the Gly-Ala repeat domain of the Epstein–Barr virus nuclear antigen 1. Proceedings of the National Academy of Sciences 94, 12616–12621 (1997).

16. M. G. Davenport, J. S. Pagano, Expression of EBNA-1 mRNA is regulated by cell cycle during Epstein-Barr virus type I latency. J Virol 73, 3154–3161 (1999).

17. H. J. Kwun et al., Kaposi’s Sarcoma-Associated Herpesvirus Latency-Associated Nuclear Antigen 1 Mimics Epstein-Barr Virus EBNA1 Immune Evasion through Central Repeat Domain Effects on Protein Processing. J Virol 81, 8225–8235 (2007).

18. R. Palich et al., Kaposi’s Sarcoma in Virally Suppressed People Living with HIV: An Emerging Condition. Cancers (Basel) 13, (2021).

19. C. Casper et al., KSHV (HHV8) vaccine: promises and potential pitfalls for a new anti-cancer vaccine. npj Vaccines 7, 108 (2022).

20. C. Li et al., Mechanisms of innate and adaptive immunity to the Pfizer-BioNTech BNT162b2 vaccine. Nature Immunology 23, 543–555 (2022).

21. H. K. Patel et al., Characterization of BNT162b2 mRNA to Evaluate Risk of Off-Target Antigen Translation. Journal of Pharmaceutical Sciences 112, 1364–1371 (2023).

22. D. S. Khoury et al., Neutralizing antibody levels are highly predictive of immune protection from symptomatic SARS-CoV-2 infection. Nature medicine 27, 1205–1211 (2021).

23. A. Muik et al., Neutralization of SARS-CoV-2 lineage B. 1.1. 7 pseudovirus by BNT162b2 vaccine–elicited human sera. Science 371, 1152–1153 (2021).

24. M. Hoffmann et al., SARS-CoV-2 variants B. 1.351 and P. 1 escape from neutralizing antibodies. Cell 184, 2384–2393. e2312 (2021).

25. J. A. Malik et al., The SARS-CoV-2 mutations versus vaccine effectiveness: New opportunities to new challenges. Journal of infection and public health 15, 228–240 (2022).

26. W. T. Harvey et al., SARS-CoV-2 variants, spike mutations and immune escape. Nature Reviews Microbiology 19, 409–424 (2021).

27. D. Planas et al., Reduced sensitivity of SARS-CoV-2 variant Delta to antibody neutralization. Nature 596, 276–280 (2021).

28. T. Braeye et al., Vaccine effectiveness against transmission of alpha, delta and omicron SARS-COV-2-infection, Belgian contact tracing, 2021–2022. Vaccine 41, 3292–3300 (2023).

29. H. S. Hillen et al., Structure of replicating SARS-CoV-2 polymerase. Nature 584, 154–156 (2020).

30. H. S. Hillen, Structure and function of SARS-CoV-2 polymerase. Current Opinion in Virology 48, 82–90 (2021).

31. K. Gangavarapu et al., Outbreak.info genomic reports: scalable and dynamic surveillance of SARS-CoV-2 variants and mutations. Nature Methods 20, 512–522 (2023).

32. A. Grifoni et al., A sequence homology and bioinformatic approach can predict candidate targets for immune responses to SARS-CoV-2. Cell host & microbe 27, 671–680. e672 (2020).

33. J. Pei, B. H. Kim, N. V. Grishin, PROMALS3D: a tool for multiple protein sequence and structure alignments. Nucleic Acids Res 36, 2295–2300 (2008).

34. D. J. Bean et al., Heterotypic immunity from prior SARS-CoV-2 infection but not COVID-19 vaccination associates with lower endemic coronavirus incidence. Sci Transl Med 16, eado7588 (2024).

35. P. A. Nesterenko et al., HLA-A∗ 02: 01 restricted T cell receptors against the highly conserved SARS-CoV-2 polymerase cross-react with human coronaviruses. Cell reports 37, (2021).

36. T. Westphal et al., Evidence for broad cross-reactivity of the SARS-CoV-2 NSP12-directed CD4+ T-cell response with pre-primed responses directed against common cold coronaviruses. Frontiers in Immunology 14, (2023).

37. E. Dolgin, T-cell vaccines could top up immunity to COVID, as variants loom large. Nat Biotechnol 40, 3–4 (2022).

38. A. Flemming, Why have T cell-inducing vaccines for HIV failed so far? Nat Rev Immunol 24, 89 (2024).

39. A. Fomsgaard, M. A. Liu, The key role of nucleic acid vaccines for one health. Viruses 13, 258 (2021).

40. C. Zhang, G. Maruggi, H. Shan, J. Li, Advances in mRNA vaccines for infectious diseases. Frontiers in immunology 10, 594 (2019).

41. M. Rechsteiner, S. W. Rogers, PEST sequences and regulation by proteolysis. Trends in biochemical sciences 21, 267–271 (1996).

42. M. Correa Marrero, I. Barrio-Hernandez, Toward understanding the biochemical determinants of protein degradation rates. ACS omega 6, 5091–5100 (2021).

43. H. J. Kwun et al., The central repeat domain 1 of Kaposi’s sarcoma-associated herpesvirus (KSHV) latency associated-nuclear antigen 1 (LANA1) prevents cis MHC class I peptide presentation. Virology 412, 357–365 (2011).

44. D. C. Tscharke et al., Identification of poxvirus CD8+ T cell determinants to enable rational design and characterization of smallpox vaccines. The Journal of experimental medicine 201, 95–104 (2005).

45. A. Varshavsky, Ubiquitin fusion technique and related methods. Methods in enzymology 399, 777–799 (2005).

46. R. T. Baker, Protein expression using ubiquitin fusion and cleavage. Current opinion in biotechnology 7, 541–546 (1996).

47. B. P. Monia, D. J. Ecker, S. T. Crooke, New perspectives on the structure and function of ubiquitin. Bio/technology 8, 209–215 (1990).

48. C. Dobaño, W. O. Rogers, K. Gowda, D. L. Doolan, Targeting antigen to MHC Class I and Class II antigen presentation pathways for malaria DNA vaccines. Immunology letters 111, 92–102 (2007).

49. M. Garzón et al., PRT6/At5g02310 encodes an Arabidopsis ubiquitin ligase of the N-end rule pathway with arginine specificity and is not the CER3 locus. FEBS Letters 581, 3189–3196 (2007).

50. A. Varshavsky, in *Cold Spring Harbor symposia on quantitative biology*. (Cold Spring Harbor Laboratory Press, 1995), vol. 60, pp. 461–478.

51. S. M. Sriram, B. Y. Kim, Y. T. Kwon, The N-end rule pathway: emerging functions and molecular principles of substrate recognition. Nature reviews Molecular cell biology 12, 735–747 (2011).

52. T. Lagousi, P. Basdeki, J. Routsias, V. Spoulou, Novel protein-based pneumococcal vaccines: assessing the use of distinct protein fragments instead of full-length proteins as vaccine antigens. Vaccines 7, 9 (2019).

53. S. K. Malladi et al., Design of a highly thermotolerant, immunogenic SARS-CoV-2 spike fragment. Journal of Biological Chemistry 296, (2021).

54. Z. Lin, S. Li, Y. Chen, Identification of viral peptide fragments for vaccine development. Methods Mol Biol 515, 261–274 (2009).

55. K. A. Hogquist et al., T cell receptor antagonist peptides induce positive selection. Cell 76, 17–27 (1994).

56. E. Cesarman et al., Kaposi sarcoma. Nature reviews. Disease primers 5, 9 (2019).

57. S. J. Gao et al., Molecular polymorphism of Kaposi’s sarcoma-associated herpesvirus (Human herpesvirus 8) latent nuclear antigen: evidence for a large repertoire of viral genotypes and dual infection with different viral genotypes. J Infect Dis 180, 1466–1476 (1999).

58. W. Meng et al., Development and characterization of a new monoclonal antibody against SARS-CoV-2 NSP12 (RdRp). Journal of medical virology 95, e28246 (2023).

59. J. T. Tellam et al., mRNA Structural constraints on EBNA1 synthesis impact on in vivo antigen presentation and early priming of CD8+ T cells. PLoS pathogens 10, e1004423 (2014).

60. G. Coppotelli, N. Mughal, D. Marescotti, M. G. Masucci, High avidity binding to DNA protects ubiquitylated substrates from proteasomal degradation. J Biol Chem 286, 19565–19575 (2011).

61. H.-G. Burgert, S. Kvist, An adenovirus type 2 glycoprotein blocks cell surface expression of human histocompatibility class I antigens. Cell 41, 987–997 (1985).

62. B. Park et al., Human cytomegalovirus inhibits tapasin-dependent peptide loading and optimization of the MHC class I peptide cargo for immune evasion. Immunity 20, 71–85 (2004).

63. J. W. Yewdell, DRiPs solidify: progress in understanding endogenous MHC class I antigen processing. Trends in immunology 32, 548–558 (2011).

64. N. Renwick et al., Multicolor microRNA FISH effectively differentiates tumor types. The Journal of clinical investigation 123, 2694–2702 (2013).

65. A. Blesch, Lentiviral and MLV based retroviral vectors for ex vivo and in vivo gene transfer. Methods 33, 164–172 (2004).

66. E. W. Sayers et al., Database resources of the national center for biotechnology information. Nucleic Acids Res 50, D20–d26 (2022).

