## Supplement for "Engineered protein destabilization reverses intrinsic immune evasion for candidate vaccine pan-strain KSHV and SARS-CoV-2 antigens"

**The PDF file includes:**

Fig. S1 to S13.

Table. S1 to S3.

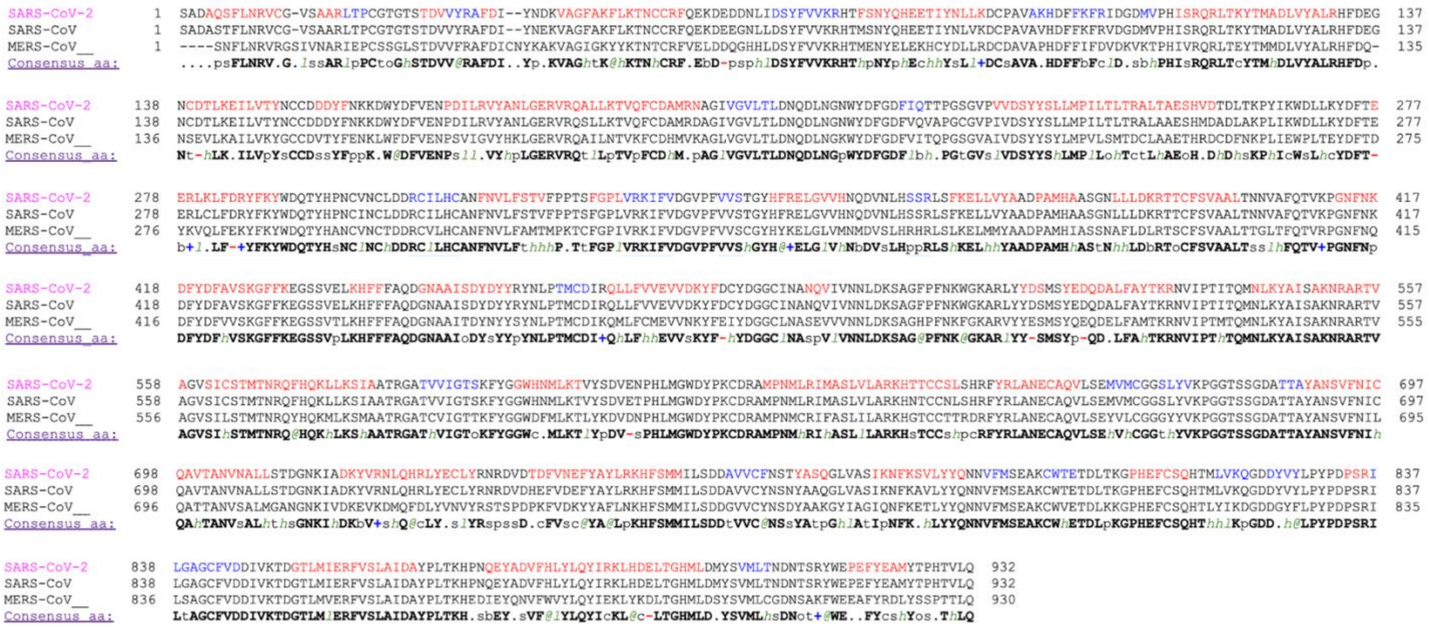

**Figure S1. Sequence alignment for SARS-CoV-2, SARS-CoV, and MERS-CoV RdRp.** Schematic is generated by PromalS3D. Red: alpha-helix, blue: beta-strand. Consensus symbols: conserved amino acids are in bold and uppercase letters; aliphatic (I, V, L): *l*; aromatic (Y, H, W, F): @; hydrophobic (W, F, Y, M, L, I, V, A, C, T, H): *h*; alcohol (S, T): o; polar residues (D, E, H, K, N, Q, R, S, T): p; tiny (A, G, C, S): t; small (A, G, C, S, V, N, D, T, P): s; bulky residues (E, F, I, K, L, M, Q, R, W, Y): b; positively charged (K, R, H): +; negatively charged (D, E): -; charged (D, E, K, R, H): c.

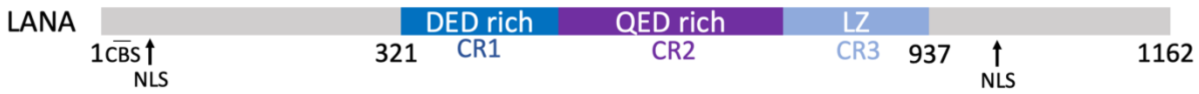


**Fig. S2. Primary structure of Kaposi sarcoma herpesvirus (KSHV) latency associated nuclear antigen (LANA).** The central repeat (CR) regions are comprised of the DED-rich CR1 (blue), QED-rich CR2 (purple), and leucine zipper CR3 (light blue) domains. N- and C-terminal portions of LANA (gray) contain unique sequences necessary for latency functions including viral episome maintenance and host chromosomal tethering: Chromatin binding sequences (CBS); nuclear localization sequences (NLS).

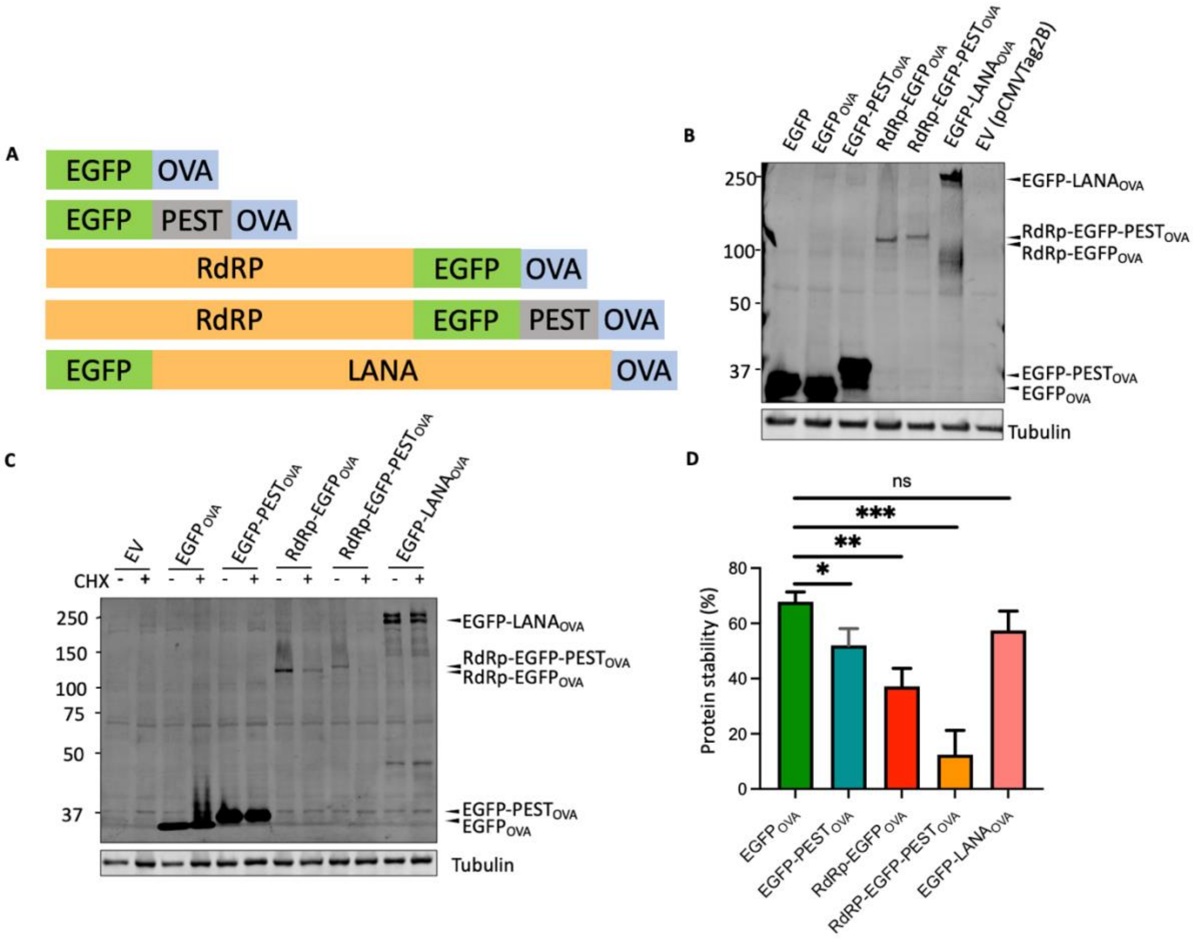


**Figure S3. SARS-CoV-2 RdRp protein is expressed at low levels, and exhibits a high turnover rate**. **(A) RdRp and LANA constructs tested for expression and stability.** Each protein was fused to EGFP and the chicken ovalbumin peptide SIINFEKL (OVA). An additional PEST sequence was also fused to RdRp to modify its protein stability. **(B) Viral proteins show low expression.** Protein expression of EGFP (lane 1), $\mathrm{EGFP_{OVA}}$,(lane 2), and even EGFP-$\mathrm{PEST_{OVA}}$ (lane 3) was robust compared to derivative proteins fused to either viral RdRp or LANA: RdRp-$\mathrm{EGFP_{OVA}}$ (lane 4), RdRp-EGFP-$\mathrm{PEST_{OVA}}$ (lane 5), EGFP-$\mathrm{LANA_{OVA}}$ (lane 6). Protein expression was determined by immunoblotting with an antibody against EGFP. **RdRp-$\mathrm{EGFP_{OVA}}$ has a high turnover rate compared to EGFP-$\mathrm{LANA_{OVA}}$.** Turnover of RdRp-$\mathrm{EGFP_{OVA}}$ and EGFP-$\mathrm{LANA_{OVA}}$ was assessed by cycloheximide (CHX) treatment 6hr prior to harvesting for **(C)** immunoblotting and **(D)** quantification by ImageJ. RdRp-$\mathrm{EGFP_{OVA}}$ (lanes 7 and 8) showed a lower stability than EGFP (lanes 3 and 4) which was further reduced by PEST fusion (lanes 9 and 10). Error bars represent SD from 3 repeats.

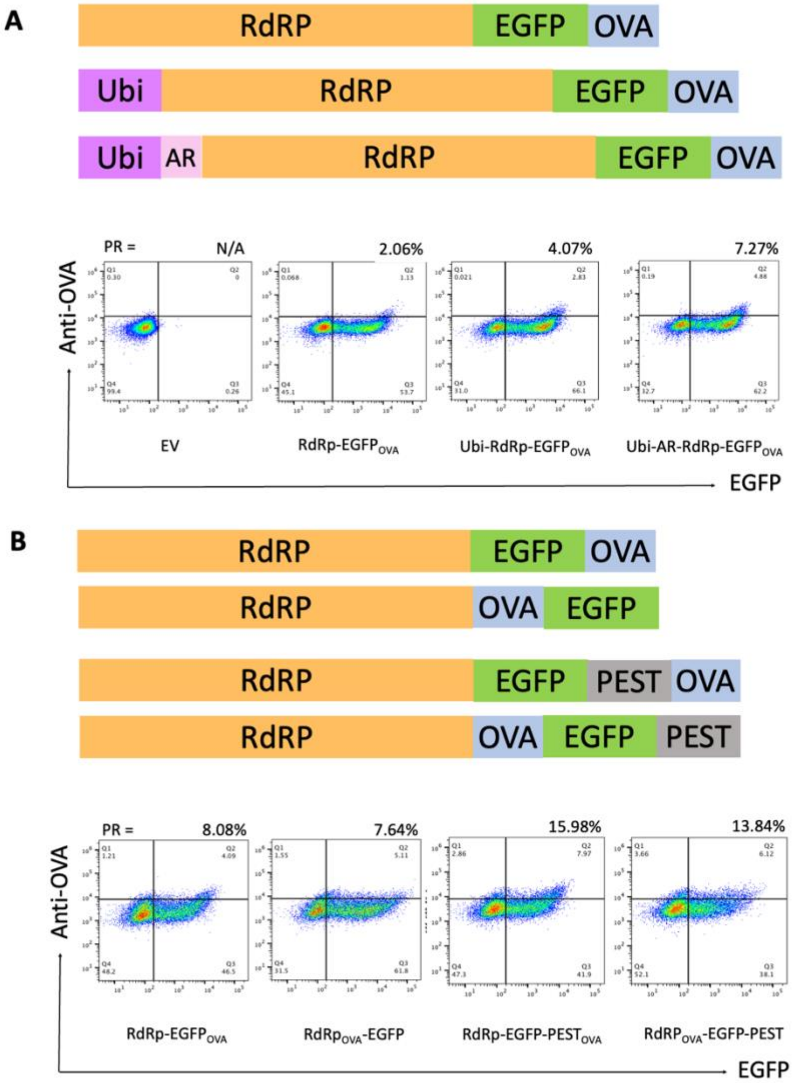


**Figure S4. Enhancement of RdRp MHC class I peptide presentation by ubiquitination and PEST fusion. (A) Ubiquitination.** Ubiquitin fused to the N-terminus of RdRp-EGFP$_{OVA}$ enhanced its presentation ratio (PR) 2-fold. Ubiquitin-AR fusion further increased PR to3.5-fold. **(B) PEST fusion.** PEST fused to RdRp-EGFP$_{OVA}$ C-terminal in orientation to OVA or C-terminal in orientation to EGFP enhanced OVA presentation 1.8 to 2-fold. Representative data derived from 3 repeats.

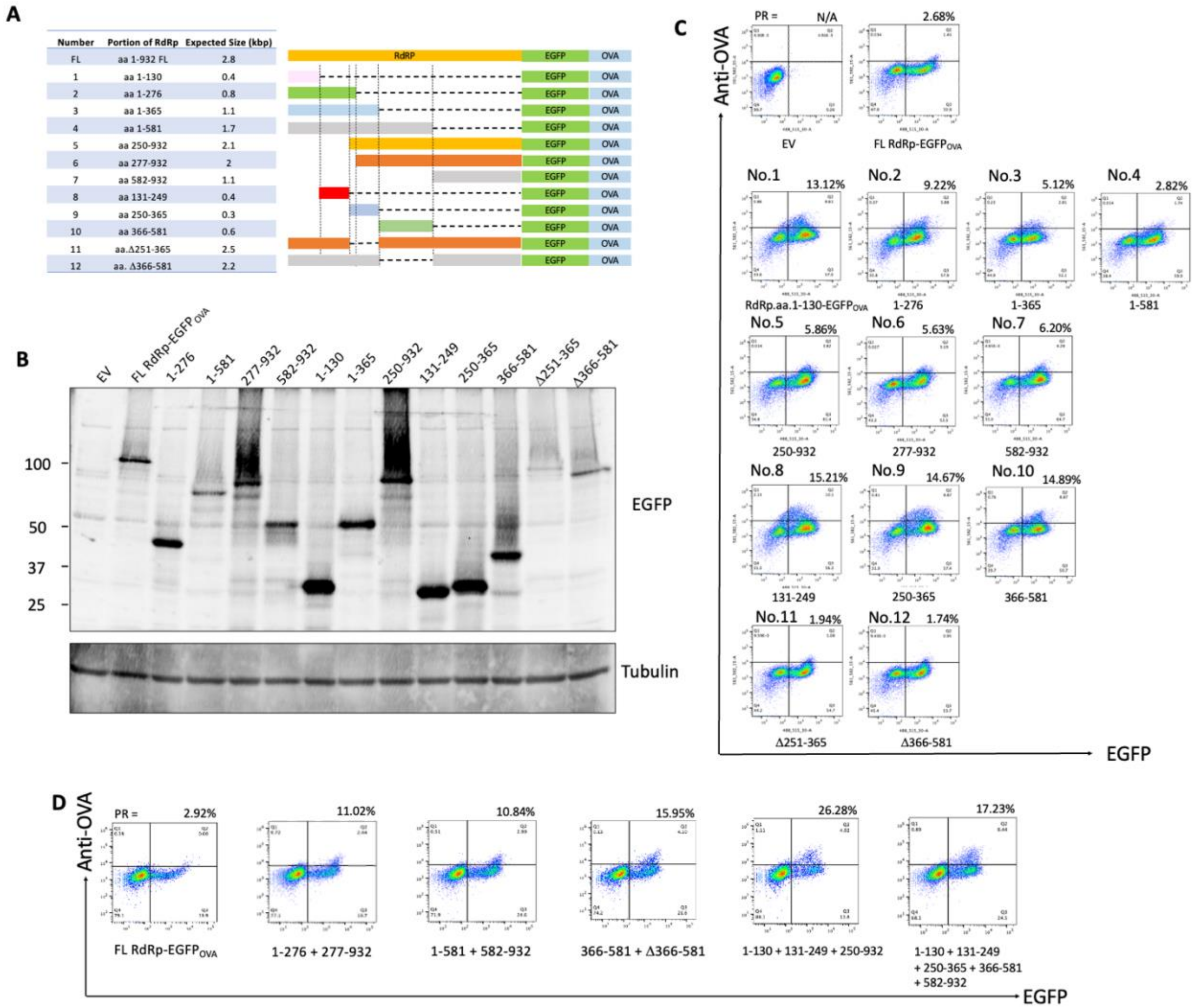


**Figure S5. RdRp MHC class I peptide presentation is enhanced by RdRp fragmentation. (A) List of fragment RdRp fragmentation constructs.** This including N-terminal fragments (No. 1-4), C-terminal fragments (No. 5-7), intermediate fragments (No. 8-10), and intermediate deletions (No. 11-12). **(B) Protein expression of fragmented RdRp-EGFP$_{OVA}$ in 293KbC2 cells.** Protein expression was validated by immunoblotting. **(C) The majority of fragmentated RdRp constructs enhanced OVA presentation.** Flow cytometry showing EGFP vs. OVA presentation of fragmented RdRp-EGFP$_{OVA}$ compared to full-length RdRp-EGFP$_{OVA}$. Representative data from 3 repeats are shown. **(D) Multiple fragmented RdRp-EGFP$_{OVA}$ in combination enhanced OVA presentation.** Flow cytometry showed an increase in PR values upon fragmentation of the full-length RdRp-EGFP$_{OVA}$ into two or more segments. Representative data from 3 replicates are presented.

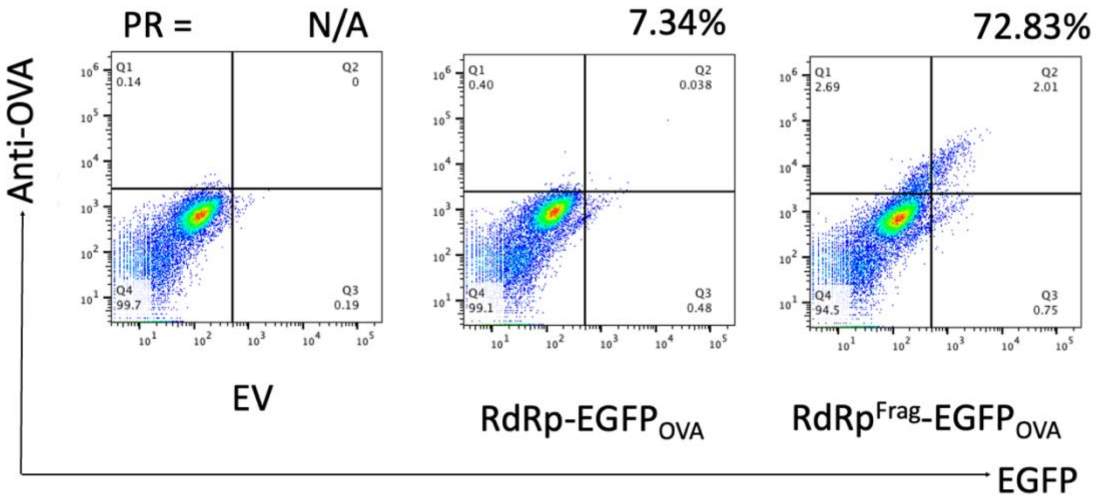


**Figure S6. Fragmented RdRp enhanced OVA presentation in the B16 melanoma cell line.** $RdRp^{Frag}$-$EGFP_{OVA}$ were transfected into B16 cells, harvested after 48h, and analyzed by flow cytometry. $RdRp^{Frag}$ demonstrated increased OVA presentation, consistent with findings in 293KbC2 cells. Representative data from a total of 3 repeats is presented.

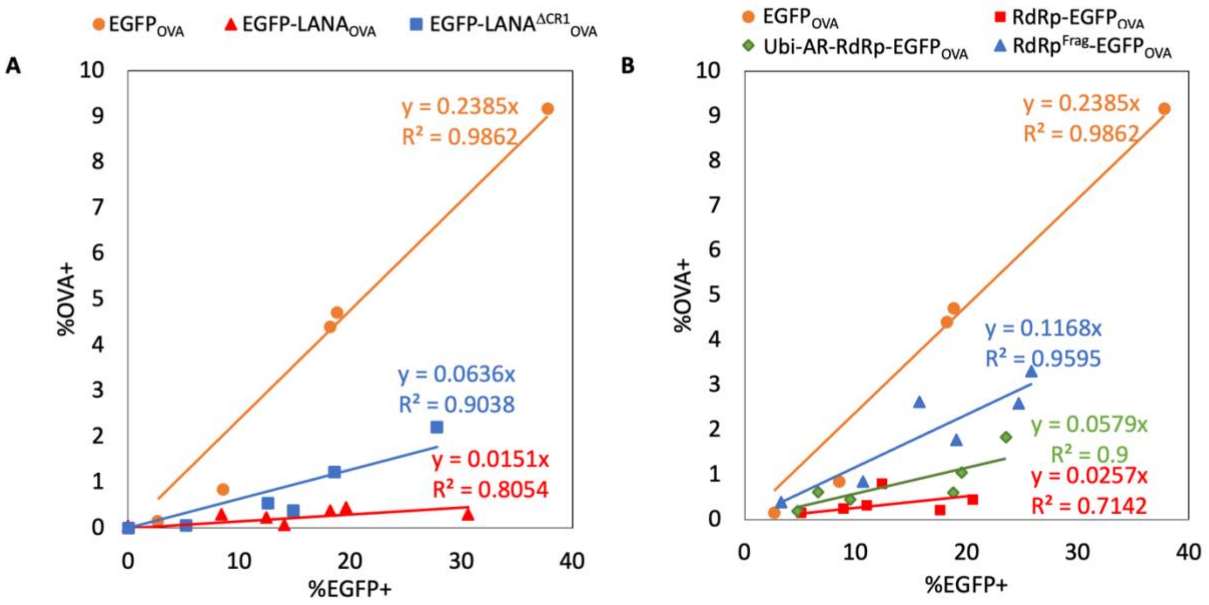


**Figure S7. Engineered LANA and RdRp constructs showed enhanced OVA presentation across varying levels of antigen expression. (A) 293KbC2 cells expressing engineered LANA proteins show enhanced OVA presentation with increasing antigen expression.** The percentage of EGFP-positive cells (Q2+Q3 in flow cytometry diagram) versus the percentage of OVA-positive cells (Q2 in flow cytometry diagram) was plotted and linearly fitted. The slope of each fitted line represents the average presentation ratio (PR) for each antigen. EGFP-LANA$^{DCR1}$$_{OVA}$ consistently enhanced OVA presentation compared to EGFP-LANA$_{OVA}$. **(B) Engineered RdRp constructs in 293KbC2 cells enhanced OVA presentation with increasing antigen expression.** Both RdRp$^{Frag}$-EGFP$_{OVA}$ and Ubi-AR-RdRp-EGFP$_{OVA}$ displayed higher OVA presentation than RdRp-EGFP$_{OVA}$, with RdRp$^{Frag}$-EGFP$_{OVA}$ exhibiting superior enhancement compared to Ubi-AR-RdRp-EGFP$_{OVA}$.

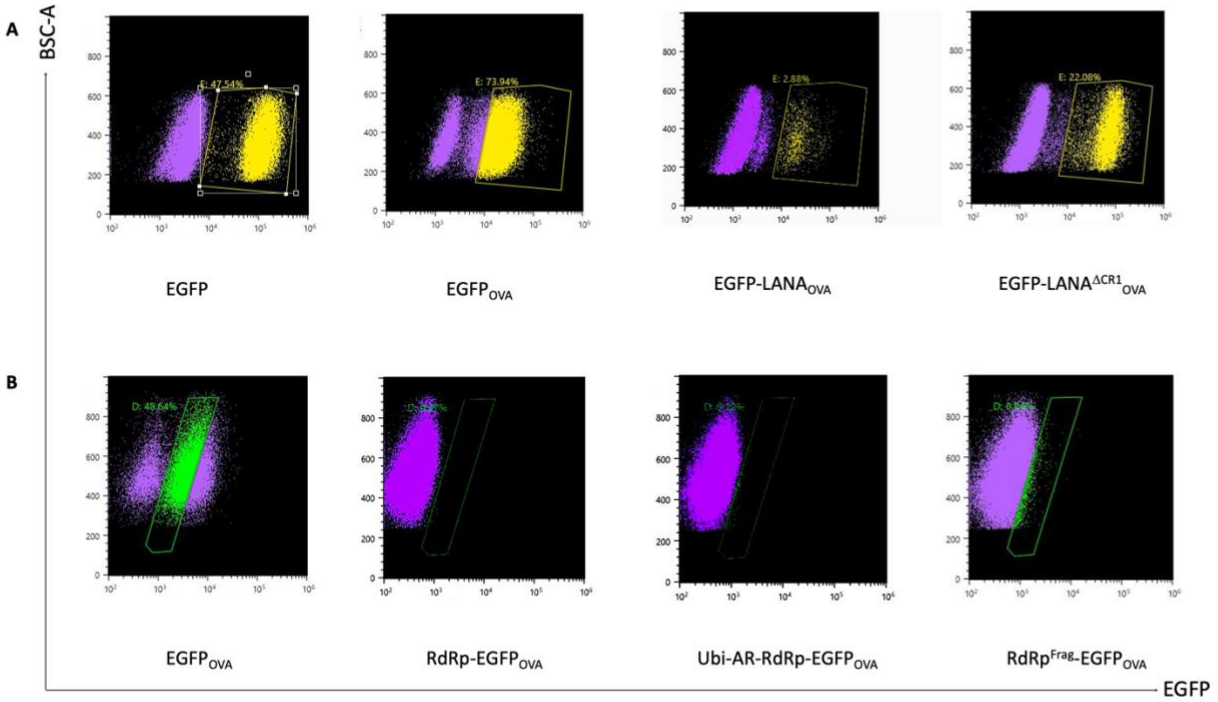


**Figure S8. MC38 Cell Sorting after lentiviral transduction to establish stable cell lines with uniform GFP expression for LANA (A) and RdRp (B) constructs.** "Yellow" gate for LANA and "green" gate for RdRp were applied. Due to the low expression level of RdRp constructs, the positive EGFP$_{OVA}$ control was gated differently than that of the LANA constructs.

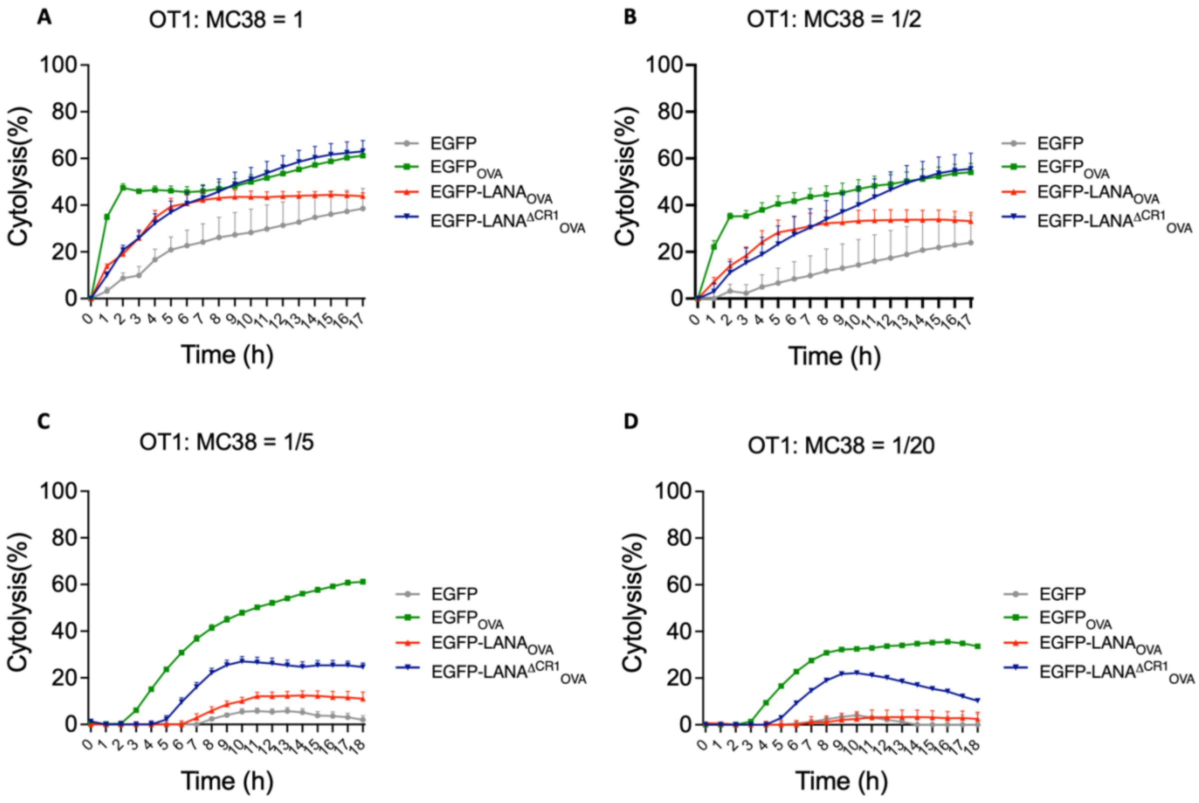


**Figure S9. Cytotoxic T lymphocyte (CTL) assay for engineered LANA at different effector: target cell ratios.** EGFP-LANA$^{DCR1}_{OVA}$ (blue line) induced more efficient OT-1 CD8 T cell killing than EGFP-LANA$_{OVA}$ (red line) at effector (OT-1): target cell (MC38) ratio= 1:1 **(A)**, 1:2 **(B)**, 1:5 **(C)**, and 1:20 **(D)**. Error bars represent SD from 4 repeats.

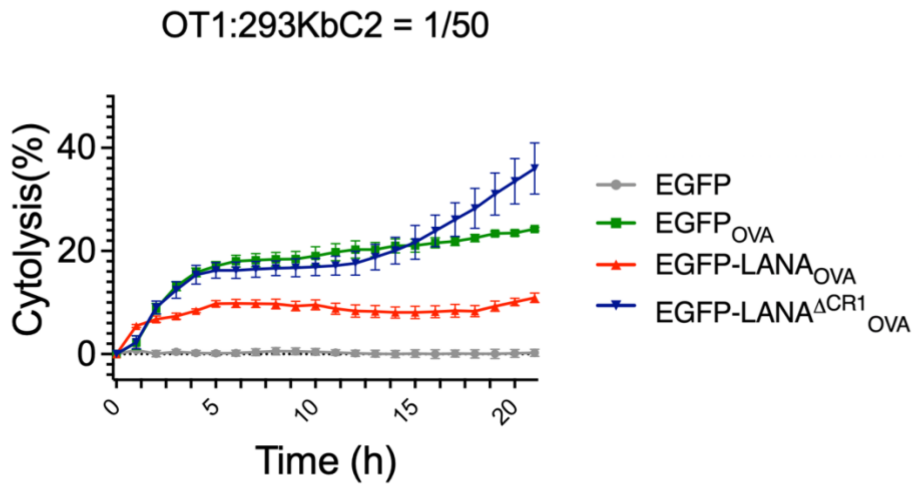


**Figure S10. Engineered LANA induced enhanced OT-1 CD8 T cell killing in 293KbC2 cells.** With transient transfection of LANA constructs into 293KbC2 cell lines, EGFP-LANA$^{DCR1}_{OVA}$ (blue line) induced OT-1 specific killing more efficiently than EGFP-LANA$_{OVA}$ (red line) Error bars represent SD from 4 repeats.

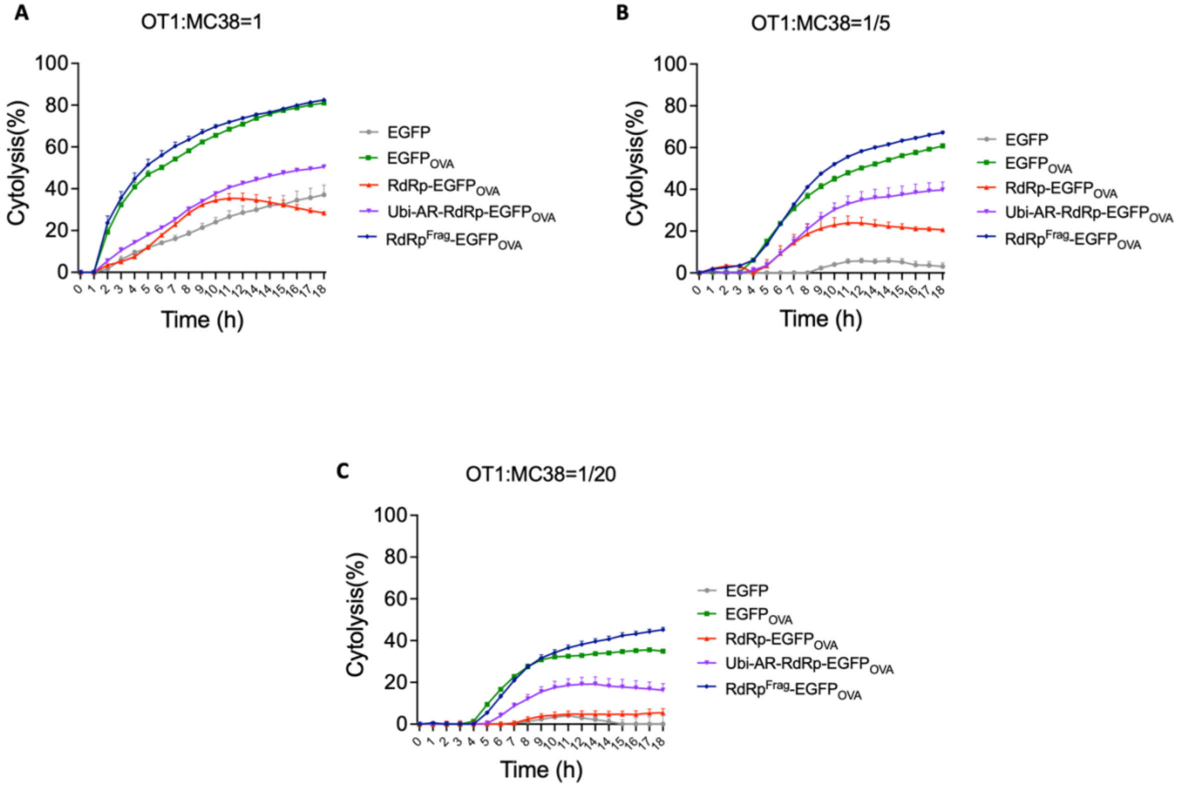


**Figure S11. Cytotoxic T lymphocyte (CTL) assay for engineered RdRp at different effector: target cell ratios.** $RdRp^{Frag}$-$EGFP_{OVA}$ (blue line) and Ubi-AR-RdRp-$EGFP_{OVA}$(mauve line) showed enhanced OT-1 CD8 T cell killing compared to RdRp-$EGFP_{OVA}$(red line) at effector (OT-1): target cell (MC38) ratio= 1:1 **(A)**, 1:5 **(B)**, and 1:20 **(C).** $RdRp^{Frag}$-$EGFP_{OVA}$ (blue line) showed better cell killing compared to Ubi-AR-RdRp-$EGFP_{OVA}$ (mauve line). Error bars represent SD from 4 repeats.

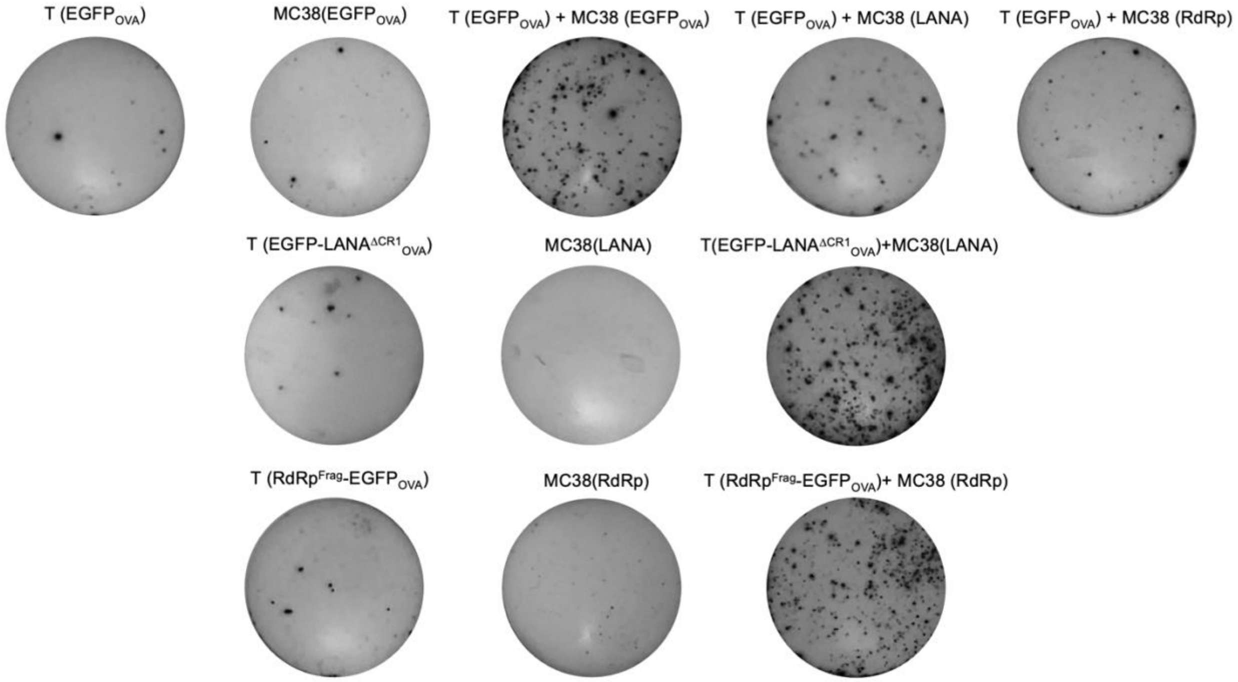


**Figure S12. Representative ELISpot wells showing the number of IFN-γ producing spots from CD8$^{+}$ T cells exposed to MC38 cells expressing wild-type viral antigens.** CD8$^{+}$ T cells from EGFP-LANA$^{\Delta CR1}$$_{OVA}$ and RdRp$^{Frag}$-EGFP$_{OVA}$ injected mice harvested 90 days post-injection were reactivated by MC38 cells expressing corresponding wild-type LANA or RdRp antigens.

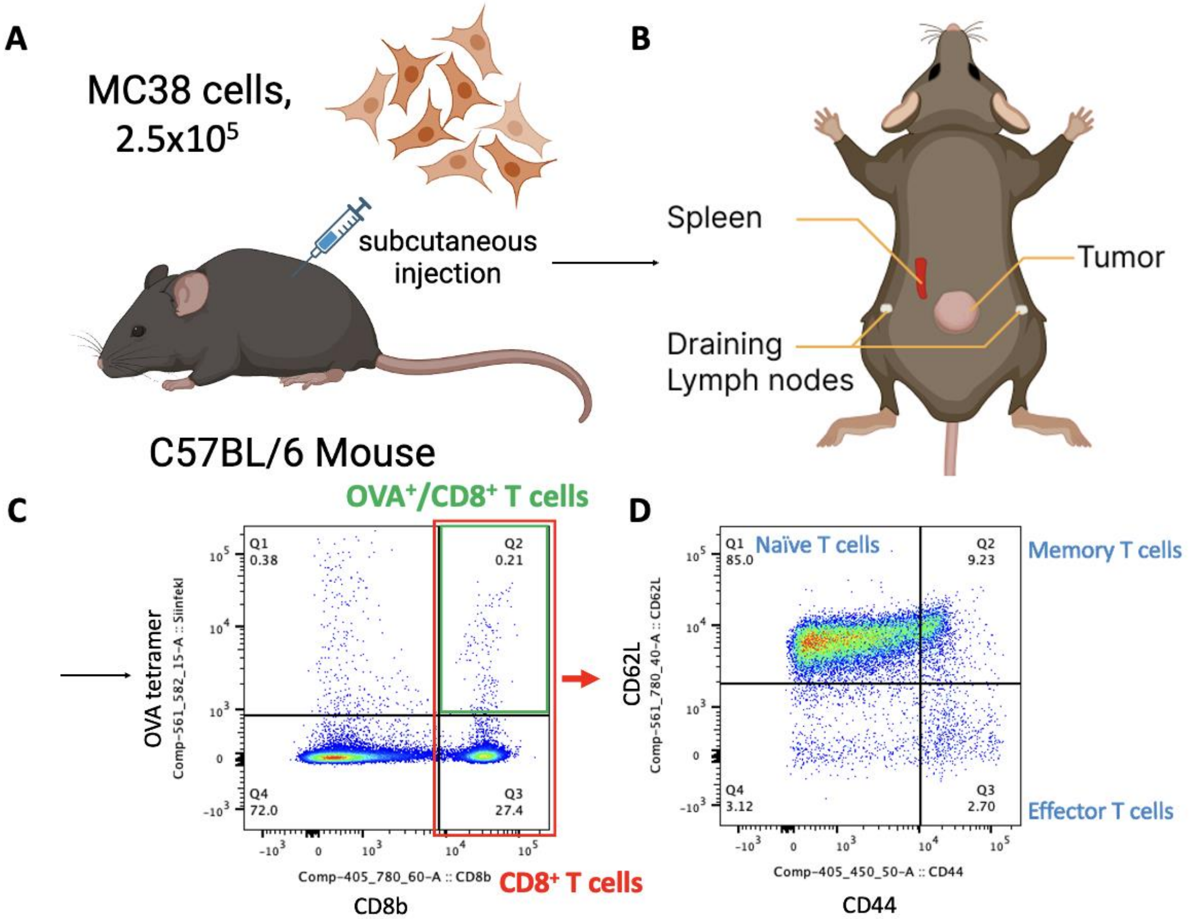


**Figure S13**. **Experimental workflow for tetramer assessment of CD8$^+$ T cell responses in Mice. (A)** C57BL/6 mice were injected subcutaneously with 2.5x10$^5$ MC38 cells. **(B)** At 7-21 days post injection, mice were sacrificed, and their tumor, draining lymph node, and spleen were harvested and stained with surface markers (CD8b, CD44, and CD62L and OVA-tetramer) for analysis. **(C)** CD8$^+$ T cell population and OVA$^+$ T cell population were determined by flow cytometry through CD8b and OVA-tetramer. **(D)** Effector and memory T cell population were determined by CD44/CD62L.

#### Table S1. Sequence for codon optimized SARS-CoV-2 RdRp

ATGTCCGCAGATGCGCAATCTTTTTTGAACAGAGTTTGTGGTGTAAGTGCAGCCCGTCTTACACCGTGCGGCACAGGCACTAGTACTGATGT
CGTCTATAGGGCGTTCGATATTTATAACGATAAAGTGGCGGGCTTCGCAAAATTCCTTAAAACTAACTGCTGTAGGTTCCAGGAAAAAGACGA
AGATGACAATCTTATAGACAGCTATTTCGTCGTTAAACGACATACTTTTAGCAACTACCAACATGAGGAAACGATCTACAACTTGCTTAAAGATT
GTCCGGCGGTAGCGAAGCACGACTTCTTCAAATTTCGGATCGATGGAGATATGGTTCCTCACATTTCCAGACAACGCCTGACTAAATACACT
ATGGCAGATCTTGTTTACGCCTTGCGACATTTTGACGAGGGTAACTGCGATACTCTTAAGGAGATCCTCGTTACCTATAATTGCTGCGATGATG
ATTACTTTAATAAGAAAGACTGGTACGATTTTGTAGAGAACCCTGATATCCTCCGGGTCTATGCGAATCTCGGGGAGCGAGTTAGACAGGCGC
TCCTTAAGACAGTGCAATTCTGCGACGCAATGCGGAATGCCGGAATAGTGGGTGTACTTACGCTTGATAATCAAGACCTCAACGGTAACTGG
TATGACTTCGGAGATTTCATCCAGACCACGCCTGGATCAGGCGTCCCCGTCGTGGATTCATATTACTCTCTGTTGATGCCTATTCTTACCCTTA
CCAGAGCTCTTACAGCTGAATCTCATGTGGATACTGATCTCACTAAACCATACATCAAGTGGGACCTCTTGAAATATGATTTTACTGAAGAAAG
GCTGAAGCTGTTCGACAGGTACTTTAAGTATTGGGACCAAACATATCATCCGAACTGTGTGAATTGTCTTGATGATCGATGCATACTCCATTGTG
CGAATTTTAATGTACTCTTCTCAACAGTCTTCCCACCTACAAGTTTTGGACCGCTCGTAAGAAAAATTTTTGTGGACGGAGTCCCTTTCGTCGT
CTCTACTGGCTATCACTTTCGAGAGTTGGGTGTCGTTCACAACCAGGACGTAAACCTTCACAGTAGTCGACTTTCCTTTAAAGAACTGCTTGT
CTACGCCGCCGACCCAGCTATGCACGCAGCCAGTGGAAACCTCTTGCTCGATAAACGAACTACCTGCTTTAGCGTTGCGGCCTTGACAAA
CAACGTCGCATTCCAGACTGTGAAGCCAGGTAATTTTAACAAAGACTTCTATGACTTTGCAGTTAGTAAGGGATTCTTTAAGGAAGGAAGTTCC
GTTGAACTCAAACATTTTTTTTTCGCGCAAGATGGCAATGCCGCTATCTCAGACTACGACTATTACCGATATAACCTGCCCACCATGTGCGAC
ATACGCCAACTGCTTTTTGTGGTCGAAGTTGTGGATAAGTACTTTGACTGCTATGATGGGGGTTGCATTAATGCAAACCAAGTCATTGTTAATAA
CCTTGATAAATCCGCTGGCTTCCCCTTTAATAAATGGGGGAAAGCACGGTTGTATTATGACTCTATGTCCTACGAGGATCAAGATGCCTTGTTT
GCGTATACCAAAAGGAACGTTATACCCACAATTACGCAGATGAATCTGAAGTATGCCATAAGTGCGAAAAATAGGGCACGCACCGTCGCAG
GAGTAAGCATATGTTCAACAATGACCAATAGGCAATTTCATCAAAAACTGTTGAAGTCAATTGCCGCCACGAGAGGCGCTACTGTCGTAATTG
GAACATCAAAATTTTATGGGGGCTGGCATAACATGCTTAAAACGGTGTATTCAGATGTCGAGAATCCGCACCTCATGGGCTGGGATTACCCG
AAGTGCGACCGGGCAATGCCGAACATGCTCAGGATCATGGCTAGTTTGGTACTGGCCCGGAAACATACCACATGCTGTTCACTCAGTCATC
GATTTTATCGACTTGCGAACGAGTGCGCCCAAGTCCTGAGCGAAATGGTAATGTGTGGAGGCTCTCTCTATGTAAAACCCGGTGGTACCTCC
TCAGGAGATGCAACTACCGCGTACGCGAACTCTGTATTTAATATTTGCCAGGCAGTAACTGCGAATGTTAATGCTCTGCTGAGCACAGATGG
GAATAAGATAGCCGATAAATACGTGCGCAACCTCCAACACCGCTTGTATGAGTGTCTGTATAGAAATCGGGACGTAGATACCGACTTTGTGAA
TGAATTTTACGCGTATCTGCGGAAGCATTTTTCTATGATGATTCTGTCCGACGATGCAGTAGTCTGCTTCAATTCAACTTATGCTTCCCAAGGCT
TGGTCGCCTCTATAAAAAATTTTAAGTCAGTCCTGTATTATCAAAATAATGTATTTATGTCAGAAGCTAAGTGTTGGACGGAGACCGATCTGACGA
AAGGTCCCCATGAGTTCTGCTCCCAACATACAATGCTGGTGAAGCAAGGTGATGATTACGTTTACCTCCCATACCCAGATCCGTCCCGGATT
CTTGGGGCTGGGTGTTTCGTGGATGATATTGTGAAAACTGATGGCACACTCATGATAGAGCGGTTTGTTTCCCTGGCTATTGATGCATACCCTC
TGACGAAGCACCCAAATCAAGAGTACGCGGACGTGTTTCACTTGTATCTGCAGTACATTCGGAAGCTGCACGACGAACTGACAGGACACAT
GCTTGATATGTACAGTGTCATGCTCACTAATGATAACACTTCCCGCTATTGGGAGCCGGAGTTTTACGAAGCGATGTATACACCCCATACTGTG
CTTCAGTAA

**Table S2. List and description of plasmid constructs. All constructs were sequence confirmed.**

| Construct | Expression | Parental vector | Chang-Moore Plasmid # | Chang-Moore Plasmid Name |
|---|---|---|---|---|
| EGFP | EGFP in mammalian cells | N/A | 2437 | pEGFP-N1 |
| EGFP-$LANA_{OVA}$ | KSHV LANA fused with EGFP and OVA peptide in mammalian cells | pCMV-Tag2B | 2452 | pCMVTag2B.eGFP.LANA1(FL).SIINFEKL |
| EGFP-$LANA^{\Delta CR1}_{OVA}$ | KSHV LANA(dCR1) fused with EGFP and OVA peptide in mammalian cells | pCMV-Tag2B | 2843 | pCMV.Tag2B.eGFP.dCR1.SIINFEKL |
| RdRp.co | Codon-optimized SARS-CoV2 RdRp in mammalian cells | pCMV14 | 4626 | p3XFLAG.CMV14-RdRp.co FIXED |
| $EGFP_{OVA}$ | EGFP fused with OVA peptide in mammalian cells | pCMV-Tag2B | 4678 | pCMV-EGFP-SIINFEKL(EcoRI) |
| PEST-$EGFP_{OVA}$ | Destablized EGFP fused with OVA peptide in mammalian cells | pCMV-Tag2B | 4677 | pCMV-d1EGFP-SIINFEKL(EcoRI) |
| RdRp-$EGFP_{OVA}$ | SARS-CoV-2 RdRp fused with EGFP and OVA peptide in mammalian cells | pCMV-Tag2B | 4681 | pCMV-Tag2B-RdRp-EGFP-SIINFEKL |
| RdRp-EGFP-$PEST_{OVA}$ | SARS-CoV-2 RdRp fused with destabilized EGFP and OVA peptide in mammalian cells | pCMV-Tag2B | 4680 | pCMV-Tag2B-RdRp-d1EGFP-SIINFEKL |
| RdRp-EGFP | SARS-CoV-2 RdRp fused with EGFP in mammalian cells | pEGFP-N1 | 4645 | pRdRp-EGFP-N1 |
| RdRp-EGFP-PEST | SARS-CoV-2 RdRp fused with destabilized EGFP in mammalian cells | pd1EGFP-N1 | 4679 | pRdRp-d1EGFP-N1 |
| $RdRp_{OVA}$-EGFP-PEST | SARS-CoV-2 RdRp fused with destabilized EGFP and OVA peptide in mammalian cells | pCMV-Tag2B | 4731 | pCMV-Tag2B-RdRp-SIINFEKL-d1EGFP |
| RdRp.1-276aa-$EGFP_{OVA}$ | Fragmented RdRp fused with EGFP and OVA peptide in mammalian cells | pCMV-Tag2B | 4690 | RdRpaa1-276-EGFP-SIINFEKL |
| RdRp.1-581aa-$EGFP_{OVA}$ | Fragmented RdRp fused with EGFP and OVA peptide in mammalian cells | pCMV-Tag2B | 4691 | RdRpaa-1-581-EGFP-SIINFEKL |
| RdRp.277-932aa-$EGFP_{OVA}$ | Fragmented RdRp fused with EGFP and OVA peptide in mammalian cells | pCMV-Tag2B | 4692 | RdRpaa-277-932-EGFP-SIINFEKL |
| RdRp.582-932aa-$EGFP_{OVA}$ | Fragmented RdRp fused with EGFP and OVA peptide in mammalian cells | pCMV-Tag2B | 4693 | RdRpaa-582-932-EGFP-SIINFEKL |
| RdRp.1-130aa-$EGFP_{OVA}$ | Fragmented RdRp fused with EGFP and OVA peptide in mammalian cells | pCMV-Tag2B | 4694 | RdRpaa-1-130-EGFP-SIINFEKL |
| RdRp.1-365aa-$EGFP_{OVA}$ | Fragmented RdRp fused with EGFP and OVA peptide in mammalian cells | pCMV-Tag2B | 4695 | RdRpaa-1-365-EGFP-SIINFEKL |
| RdRp.250-932aa-$EGFP_{OVA}$ | Fragmented RdRp fused with EGFP and OVA peptide in mammalian cells | pCMV-Tag2B | 4696 | RdRpaa-250-932-EGFP-SIINFEKL |
| RdRp.131-249aa-$EGFP_{OVA}$ | Fragmented RdRp fused with EGFP and OVA peptide in mammalian cells | pCMV-Tag2B | 4697 | RdRpaa-131-249-EGFP-SIINFEKL |
| RdRp.250-365aa-$EGFP_{OVA}$ | Fragmented RdRp fused with EGFP and OVA peptide in mammalian cells | pCMV-Tag2B | 4698 | RdRpaa-250-365-EGFP-SIINFEKL |
| RdRp.366-581aa-$EGFP_{OVA}$ | Fragmented RdRp fused with EGFP and OVA peptide in mammalian cells | pCMV-Tag2B | 4699 | RdRpaa-366-581-EGFP-SIINFEKL |
| RdRp.Δ251-365aa-$EGFP_{OVA}$ | Fragmented RdRp fused with EGFP and OVA peptide in mammalian cells | pCMV-Tag2B | 4710 | RdRpaa-del.251-365-EGFP-SIINFEKL |
| RdRp.Δ366-581aa-$EGFP_{OVA}$ | Fragmented RdRp fused with EGFP and OVA peptide in mammalian cells | pCMV-Tag2B | 4711 | RdRpaa-del.366-581-EGFP-SIINFEKL |
| Ubi-$EGFP_{OVA}$ | N-terminus ubiqutin fused EGFP-SIINFEKL in mammalian cells | pCMV-Tag2B | 4707 | Ubiquitin-EGFP-SIINFEKL |
| Ubi-RdRp-$EGFP_{OVA}$ | N-terminus ubiqutin fused RdRp-EGFP-SIINFEKL in mammalian cells | pCMV-Tag2B | 4708 | Ubiquitin-RdRp-EGFP-SIINFEKL |
| Ubi-AR-$EGFP_{OVA}$ | Modified N-terminus ubiqutin fused EGFP-SIINFEKL in mammalian cells | pCMV-Tag2B | 4715 | Ubiquitin(optimized)-EGFP-SIINFEKL |
| Ubi-AR-RdRp-$EGFP_{OVA}$ | Modified N-terminus ubiqutin fused RdRp-EGFP-SIINFEKL in mammalian cells | pCMV-Tag2B | 4716 | Ubiquitin(optimized)-RdRp-EGFP-SIINFEKL |
| pLenti-$EGFP_{OVA}$ | Lentiviral transduction of EGFP-SIINFEKL to MC38 cells | pLVX EF.Puro | 4769 | pLenti-Puro-EGFP-SIINFEKL |
| pLenti-$RdRp_{OVA}$-EGFP-PEST | Lentiviral transduction of RdRp-SIINFEKL-EGFP-PEST to MC38 cells | pLVX EF.Puro | 4770 | pLenti-Puro-RdRp-SIINFEKL-d1EGFP |
| pLenti-RdRp-$EGFP_{OVA}$ | Lentiviral transduction of RdRp-EGFP-SIINFEKL to MC38 cells | pLVX EF.Puro | 4771 | pLenti-Puro-RdRp-EGFP-SIINFEKL |
| pLenti-Ubi-AR-RdRp-$EGFP_{OVA}$ | Lentiviral transduction of ubiquitin-A-R-RdRp-EGFP-SIINFEKL to MC38 cells | pLVX EF.Puro | 4772 | pLenti-Puro-Ubiquitin-RdRp-EGFP-SIINFEKL |
| pLenti-RdRp366-581aa-$EGFP_{OVA}$ | Lentiviral transduction of RdRp.366-581-EGFP-SIINFEKL to MC38 cells | pLVX EF.Puro | 4773 | pLenti-Puro-RdRp366-581-EGFP-SIINFEKL |
| pLenti-RdRpΔ366-581aa-$EGFP_{OVA}$ | Lentiviral transduction of RdRp.del.366-581-EGFP-SIINFEKL to MC38 cells | pMuLE EF.Bla | 4779 | pLenti-RdRp.del.366-581-EGFP-SIINFEKL |
| pLenti-EGFP-$LANA_{OVA}$ | Lentiviral transduction of EGFP-LANA-SIINFEKL to MC38 cells | pLVX EF.Puro | 4778 | pLenti-Puro-EGFP-LANA1-SIINFEKL |
| pLenti-EGFP-$LANA^{\Delta CR1}{}_{OVA}$ | Lentiviral transduction of EGFP-LANA(dCR1)-SIINFEKL to MC38 cells | pLVX EF.Puro | 4876 | pLenti-Puro-EGFP-LANA1dCR1-SIINFEKL |
| psPAX2 | Packaging plasmid for lentiviral transduction | N/A | 2504 | psPAX2 |
| pMD2.G | Packaging plasmid for lentiviral transduction | N/A | 2505 | pMD2.G |

### Table S3. List of primers for plasmid construction

| Constructs | Chang-Moore Plasmid # | Primer-Sense | Primer- Antisense | PCR Template |
|---|---|---|---|---|
| RdRp-EGFP | 4645 | GCACGAATTCAACATGTCCGCAGATGCGCAATC | AGAGGATCCACCTGAAGCACAGTATGGGGTG | RdRp.co |
| RdRp-EGFP-PEST | 4679 | GCACGAATTCAACATGTCCGCAGATGCGCAATC | AGAGGATCCACCTGAAGCACAGTATGGGGTG | RdRp.co |
| RdRp.Δ366-581aa-EGFP$_{OVA}$ | 4711 | GTAGCGCCTCTCGTTCGACTACTGTGAAGGTT | AACCTTCACAGTAGTCGAACGAGAGGCGCTAC | RdRp-EGFP$_{OVA}$ |
| Ubi-EGFP$_{OVA}$ | 4707 | TATCGCCGCGGATGCAGATCTTCGTG | TAGCGAATTCAGCCTAGCACCTCTGAGACGGAG | pRK5-HA-Ubiquitin-WT (Addgene #17608) |
| Ubi-RdRp-EGFP$_{OVA}$ | 4708 | TATCGCCGCGGATGCAGATCTTCGTG | TAGCGAATTCCCTAGCACCTCTGAGACGGAG | |
| Ubi-AR-EGFP$_{OVA}$ | 4715 | ATCGCCGCGGATGCAGATCTTCGTG | TAGCGAATTCAGCCTACG AGCTCTGAGACGGAG | |
| Ubi-AR-RdRp-EGFP$_{OVA}$ | 4716 | ATCGCCGCGGATGCAGATCTTCGTG | TAGCGAATTCCCTACGAGCTCTGAGACGGAG | |
| pLenti-RdRpΔ366-581aa-EGFP$_{OVA}$ | 4779 | ATCGAGCGCTAATTAACCCCTCACTAAAGGG | ATCGTCTAGATAATTAAGGTACCGGGC | RdRp.Δ366-581aa-EGFP$_{OVA}$ |
| pLenti-EGFP$_{OVA}$ | 4769 | ATCGAGCGCTAATTAACCCCTCACTAAAGGG | ATCGCCTGCAGGTAATTAAGGTACCGGGC | EGFP$_{OVA}$ |
| pLenti-RdRp$_{OVA}$-EGFP-PEST | 4770 | | | RdRp$_{OVA}$-EGFP-PEST |
| pLenti-RdRp-EGFP$_{OVA}$ | 4771 | | | RdRp-EGFP$_{OVA}$ |
| pLenti-Ubi-AR-RdRp-EGFP$_{OVA}$ | 4772 | | | Ubi-AR-RdRp-EGFP$_{OVA}$ |
| pLenti-RdRp366-581aa-EGFP$_{OVA}$ | 4773 | | | RdRp366-581aa-EGFP$_{OVA}$ |
| pLenti-EGFP-LANA$_{OVA}$ | 4778 | | | EGFP-LANA$_{OVA}$ |
| pLenti-EGFP-LANA$^{\Delta CR1}{}_{OVA}$ | 4876 | | | EGFP-LANA$^{\Delta CR1}{}_{OVA}$ |
